# SARS-CoV-2 envelope-protein corruption of homeostatic signaling mechanisms in mammalian cells

**DOI:** 10.1101/2021.06.16.448640

**Authors:** Tobias Schulze, Andreas Hartel, Sebastian Höler, Clara Hemming, Robert Lehn, Dominique Tandl, Timo Greiner, Adam Bertl, Kenneth Shepard, Anna Moroni, Gerhard Thiel, Oliver Rauh

## Abstract

During a SARS-CoV2 infection, host cells produce large amounts of the viral envelope protein (Ep-CoV2). Ep-CoV2 is partially inserted into the membrane of nascent viral particles and into cellular membranes. To mimic the pathophysiological impact of the cellular protein fraction, Ep-CoV2 was overexpressed in mammalian cells and effects on key signaling parameters were monitored. By tagging with green fluorescent protein (GFP), we found that Ep-CoV2 protein is mostly present in the endoplasmic reticulum with additional trace amounts in the plasma membrane. We observed that wild-type Ep-CoV2 and, to a lesser extent, its mutants (N15A, V25F) corrupted some of the most important homeostatic mechanisms in cells. The same was observed with isolated transmembrane domains of the protein. The Ep-CoV2-evoked elevation of intracellular Ca^2+^ and pH as well as the induced membrane depolarization produced by the presence of the protein interfere with major signal transduction cascades in host cells. These functions of Ep-CoV2, which likely contribute to the pathogenesis of the viral protein, result from the ion-channel activity of the viral protein. Two independent assays, a functional reconstitution of Ep-CoV2 protein in artificial membranes and a rescue of K^+^-deficient yeast mutants, confirm that Ep-CoV2 generates a cation-conducting channel with a low unitary conductance and a complex ion selectivity. The data presented here suggest that specific channel function inhibitors of Ep-CoV2 can provide cell protection and virostatic effects.

## INTRODUCTION

Viral infections with different strains of the coronavirus (CoV) have led in the last decades to the severe acute respiratory syndrome (SARS)-CoV1 (2003) and Middle East respiratory syndrome (MERS) (2012) epidemics and to the most recent COVID-19 (Coronavirus disease 2019) pandemic (2019). CoVs are positive-strand enveloped RNA viruses with many different hosts and strains. The alpha-coronavirus strains 229E and OC43 are associated with up to 30% of common cold cases showing mild symptoms in humans [1], while in non-human strains, such as the bovine corona virus (beta-coronavirus) or the avian infectious bronchitis virus (IBV, gamma-coronavirus), infections have large impacts in agricultural settings [2].

CoVs are built from four major structural proteins: the nucleocapsid (N) protein engulfing the viral RNA, the spike (S) protein, the membrane (M) protein and the envelope (E) protein. The S-, M- and E-proteins are membrane proteins embedded in a lipid bilayer surrounding the ribonucleoprotein complex in the virus particle [3]. Presently, academic and pharmaceutical efforts are mainly focused on the S-protein which is involved in the entry of the virus into the host cell [4]. The M- and E-proteins in contrast are less well studied [3] but there is compelling evidence for a concerted role of both proteins in the formation of virus-like particles in the ER-Golgi intermediate compartment (ERGIC) as well as in the process of budding of the mature virus particles from the host [5]. A prominent functional role of the E-protein is further supported by *in vivo* studies, which show that its genetic depletion alone leads to a marked decrease of viral fitness leading to reduced disease [6,7]. These data together with the indirect evidence for a functional role of the E-protein in acute respiratory distress syndrome [3] underscore the necessity for improving our understanding of the role of the E-protein in host-virus interaction.

E-proteins are composed of three structural domains – a short hydrophilic N-terminus, a long transmembrane domain, and a long hydrophilic C-terminal domain with an alpha helical fold [8]. Computer-based models and structural studies on purified proteins in detergent micelles suggest that the E-protein from both SARS-CoV1 and SARS-CoV2 forms a pentamer with a central pore that could serve as an ion channel [8-11].

For an understanding of the functional role of the E-protein in viral infection and replication, a detailed characterization of ion channel activity is essential. In the case of the E-protein from CoVs a number of electrophysiological studies have been performed using purified or synthetic E-proteins reconstituted in planar lipid bilayers [12-18]. In addition, voltage clamp measurements were carried out in mammalian cells transiently transfected with the viral protein [19]. The general consensus from all these studies is that the viral protein indeed exhibits channel activity. However, detailed scrutiny of the data provides a rather diverse picture on basic functional features of this channel, including unitary conductance and ion selectivity. While some studies report, after functional reconstitution of the protein in planar lipid bilayers, distinct channel fluctuations with a conductance in the lower pS range (10-20 pS) [18,20], others find a wide spectrum of conductance including unitary conductance in the range from 19 to >400 pS [12-14]. While all studies agree on an overall cation selectivity of the channel, there is no consensus on its preference for K^+^ over Na^+^ [21] and whether it is at all permeable to Ca^2+^ [17, 21]. In some of these studies it was reported that E-protein-mediated channel fluctuations were inhibited by the antiviral channel blocker amantadine or hexamethylene-amiloride (HMA), and the presence of these compounds also resulted in an attenuation of virus replication *in vivo* [11,19, 22, 23]. Because of the heterogeneity in the functional data, it is currently difficult to interpret these pharmacological results in the context of a defined channel activity of the E-protein. Accordingly, it is also not surprising that a correlation between the sensitivity of the channel to HMA and its effect on plaque formation has been questioned [21].

Furthermore, reports on whether the E-protein from SARS-CoV1 (Ep-CoV1) and SARS-CoV2 (Ep-CoV2) can form ion channels that reach the plasma membrane of cells are controversial. While some studies report an Ep-CoV1-generated conductance in the plasma membrane of *Xenopus* oocytes and mammalian cells [19,24], others do not find any evidence for elevated currents after expressing the protein in the same systems [25,26]. The latter work suggests that the presence or absence of the Ep-CoV2 in the plasma membrane is a matter of protein sorting. Only after addition of an appropriate sorting signal trafficking of Ep-CoV2 was reported to the plasma membrane of *Xenopus* oocytes and mammalian cells where it generated a cation-selective conductance [25]. At this point, it is difficult to explain this variability in observed channel function. One possible explanation is that this functional diversity reflects differences in the experimental conditions, including the systems under which the E-protein was isolated or synthesized and the approaches with which it was functionally reconstituted.

To further improve understanding of the functional properties of Ep-CoV2, we first confirmed the channel function of Ep-CoV2. When the protein was synthesized *in vitro* into nanodiscs and reconstituted into model bilayer membranes, it generated a cation conductance with a defined low unitary amplitude and long open and closed times on the order of seconds. Heterologous expression of the protein and its mutants in mammalian cells caused major changes of cellular signaling parameters, specifically membrane depolarization, an elevation of free intracellular Ca^2+^ concentration ([Ca^2+^]_in_) and an intracellular alkalization. It is already well established that a number of viral membrane proteins exhibit channel activity with well-known roles in distinct steps of viral entry, replication and budding [27-29]. Particularly interesting in this context is the ability of some of these channels to modulate the concentration of free Ca^2+^ in the cytosol of the host cells [30]. The latter is a crucial second messenger in mammalian cells which orchestrates a multitude of cellular functions. A modulation of these signaling cascades is a key event in the infection and replication strategy of many viruses [30-32]. Hence, like in the case of other viral membrane channel proteins, such as the M2 Protein from Influenza A [33], the E-protein is a good target for the development of blockers that may act as virostatic drugs.

## RESULTS

### The E-protein of SARS-CoV2 forms functional ion channels in model membranes

Several studies have reported that the E-protein from SARS-CoV1 (Ep-CoV1) forms an ion channel [3]. The high similarity in the primary amino acid sequence with the corresponding Ep-CoV2 from SARS-CoV2 (Fig. 1A,B) suggests that the latter should also exhibit channel function, which has been recently confirmed[18, 20]. We also observed this function in experiments in which Ep-CoV2 was synthesized *in vitro* into membrane-mimetic nanodiscs, a method with a low propensity of channel contaminations from the expression system [34]. By using biochemical methods, we show that nanodiscs after cell-free synthesis contain the FLAG-tagged Ep-CoV2 (Ep-CoV2_FLAG_) at different oligomeric states (Figure 1C), confirming the propensity of the protein to spontaneously form multimers in membranes [26]. We have not specified the full range of possible oligomeric states in the nanodiscs and our assay does not exclude the presence of pentamers, the presumed channel forming structure [8,10], in this system. After functional reconstitution of these EP-CoV2_FLAG_ proteins from the nanodiscs into a planar lipid bilayer (1:1 mixture of DPhPC:DPhPS), distinct channel fluctuations with characteristic seconds-long open and closed states and a low unitary amplitude became visible (Fig. 1D,E). Typically, this type of activity was observable 10-20 minutes after the nanodisc/protein complexes were added to the model membrane. For FLAG-tagged and untagged Ep-CoV2, we found, in symmetrical solutions with 250 mM KCl, a dominant unitary conductance of 16.3 ± 0.3pS and 18.0 ± 0.4pS, respectively. Hence addition of the FLAG-tag at the amino terminus of the protein has no major effect on the channel conductance (Fig. 1E). The i/V relation of the channel is linear over the range of test voltage between ±150 mV. The unitary conductance increases with increasing salt concentration; a 10-fold elevation in KCl from 0.1 to 1 M increases the unitary channel conductance by a factor of 2.3 (Supplementary Figure 1A, B) The open-channel conductance is sensitive to the lipid bilayer composition. In a bilayer with a 1:1 mix of neutral (DPhPC) and anionic lipids (DPhPS) the unitary conductance increased 1.5-fold over that recorded in neutral bilayers (Supplementary Figure .1A, B). A sensitivity of Ep-CoV1 channel conductance to anionic phospholipids was already reported previously [15].

**Figure 1.**
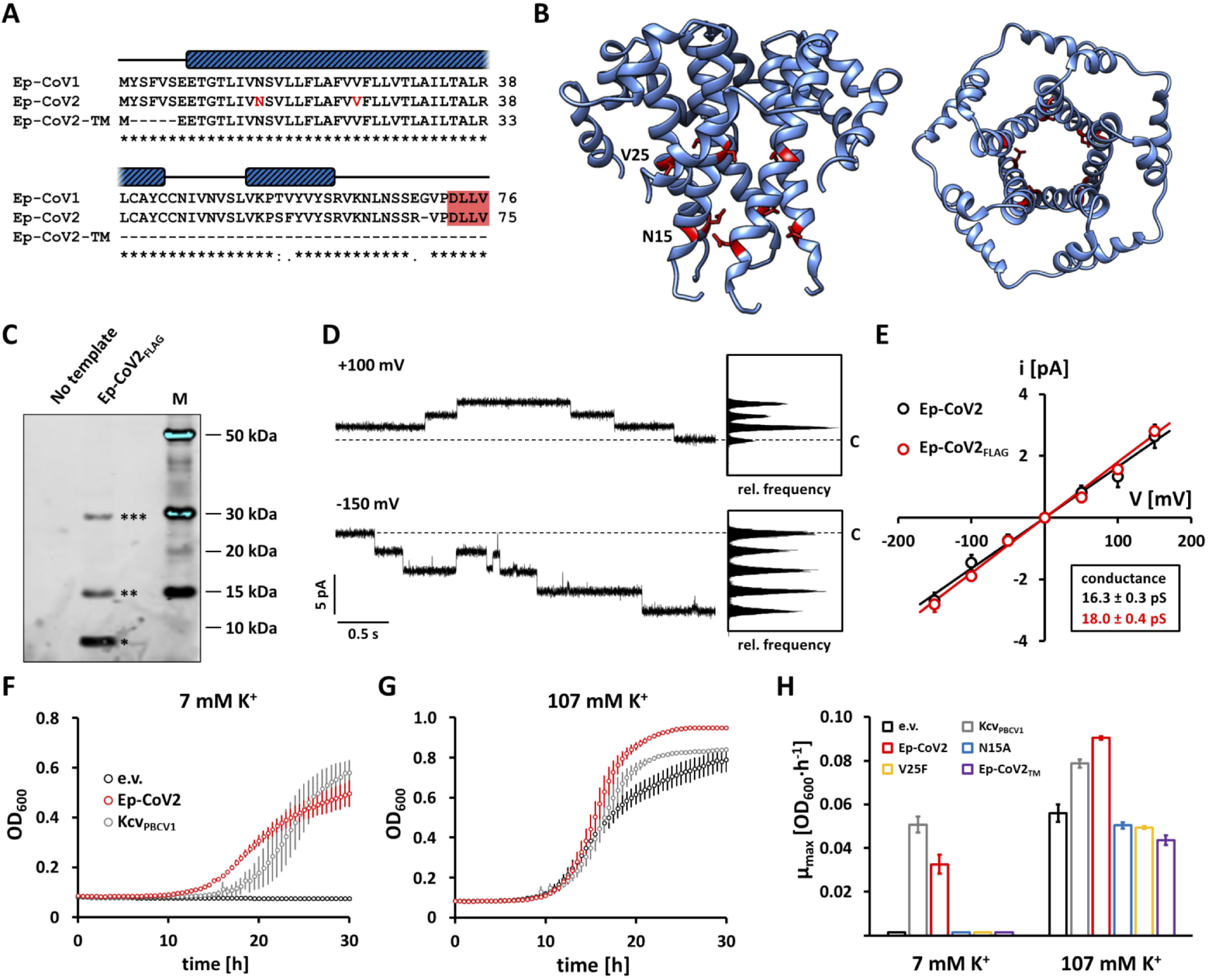
Ep-CoV2 shows ion-channel activity in artificial lipid bilayers and yeast-complementation assay. **(A)** Alignment of the amino acid sequences of envelope proteins from SARS-CoV1 (Ep-CoV1), SARS-CoV2 (Ep-CoV2) and the truncated version of Ep-CoV2 (Ep-CoV2-TM) used in this study. Blue bars above sequences indicate regions with α-helical structure according to NMR analysis of full-length SARS-CoV2 [11]. Asterisks denote conserved amino acids in Ep-CoV1 and Ep-CoV2; colons and dots indicate conservative and semi-conservative amino acid exchanges, respectively. Amino acids N15 and V25, which were mutated in this study, are colored red. The PDZ-binding motif D-L-L-V at the C-terminus of Ep-CoV1 and Ep-CoV2 highlighted by red box. **(B)** Cartoon representation of homology model of Ep-CoV2 (amino acids 8 to 65). Left: side view of pentameric channel. Right: view from top to bottom. Positions of amino acids N15 and V25 highlighted in red. Homology model created with Swiss-Model using NMR structure of SARS Coronavirus E-protein (PDB 5×29) as template. **(C)** Western blot analysis of purified No-template and Ep-CoV2_FLAG_ *in vitro* expression products using a mouse anti-FLAG tag antibody as described in material and methods. Number of asterisks indicates the oligomeric state of Ep-CoV2 _FLAG_ assuming a monomer weight of ca. 9 kDa. **(D)** Exemplary current fluctuations generated by *in vitro* translated Ep-CoV2 channels in 1:1 DPhPC/DPhPS lipid bilayer at +100 and -150 mV in symmetrical 250 mM KCl, 1 mM EGTA, and 10 mM HEPES (pH 7.4); corresponding amplitude histograms are depicted on the right. Dashed lines indicate the zero-current level. Current traces filtered at 100 Hz for visualization. **(E)** Unitary open-channel currents of Ep-CoV2 and Ep-CoV2_FLAG_. Solid lines are best fits of i = γ·V to the two data sets, resulting in unitary conductance γ for Ep-CoV2 (black) and Ep-CoV2_FLAG_ (red) given in box on bottom right. **(F-H)** Results of functional yeast complementation assay as described in material and methods using yeast strain PLY240 which is lacking K_+_ -uptake systems Trk1 and Trk2. Growth of PLY240 cells transformed with empty vector (e.v.), Ep-CoV2 or Kcv_PBCV1_ was measured in low (7 mM) K_+_ **(F)** and high (107 mM) K_+_ **(G). (H)** Maximal growth rates µ_max_ calculated from growth curves as in (F) and (G). Data points in **E-H** are arithmetic means ± standard deviations of >3 independent measurements.

Altogether the channel recordings show some similarities to published data in that the unitary conductance is in the range of 10 to 20 pS, values consistent with those that have been reported by others [11,12, 18,20]. Differing from some previous studies [11, 12], however, the channel in our recordings has only one major conductance. Worth noting are the longer open and closed dwell times in our recordings compared to those reported by Hutchison et al. [20]. In both studies, the viral proteins were solubilized either into nanodiscs (the present study) or into amphipols [20] before functional reconstitution in bilayers. It is possible that this preparative step, which may preserve the native fold of the oligomeric protein, is the source of these differences in the functional properties of the channel.

Control experiments confirm that the channel activity measured here is indeed generated by Ep-CoV2. In experiments in which *in vitro* expression was performed as in Fig. 1D but without the vector or without the nanodiscs we never detected any channel activity in the bilayer. As an additional control, the Ep-CoV2 N15A mutant was synthesized into nanodiscs using the same methods employed for the wt-protein. This mutant, which is expected to corrupt channel function [14], was in our experiments able to generate some channel activity. However, this occurred at a lower efficiency than with the wt-channel. While channel activity of the Ep-CoV2-wt was robustly seen in typical single-channel recordings after 10-20 minutes, the N15A mutant, at comparable protein concentration, showed only activity after 240 minutes (three recordings in an accumulated experimental time of 12-14 hrs) (Supplementary Figure 1C). The results of these experiments indicate that the N15A mutant is, in principle, active, but the mutant has either a reduced ability to form functional ion channels or the mutation compromises the pore stability and thus causes considerably reduced channel open probabilities.

Channel function of Ep-CoV2 was further confirmed by an independent yeast complementation assay [35]. A yeast mutant lacking the major K^+^ uptake systems Trk1 and Trk2 grows in medium with high (107 mM) K^+^, independent of whether the cells express the viral K^+^ channel Kcv_PBCV1_ or Ep-CoV 2. This confirms that the latter protein is not cytotoxic for yeast growth. The data in Figs. 1F-H show that this mutant strain, however, is unable to grow in a medium with a low (7 mM) K^+^ concentration. This growth defect can be rescued by expressing the small K^+^ channel Kcv_PBCV1_ as a positive control [36]. A similar growth rescue was achieved by expressing Ep-CoV2 (Fig. 1F,H). These data are in agreement with the results from bilayer recordings in that Ep-CoV2 forms a channel with a K^+^ conductance. The finding that expression of Ep-CoV2 is able to compensate for the absence of an endogenous K^+^ uptake system is also in agreement with similar experiments in which the expression of the viral protein was modulating the growth of bacteria [37], both indicating that Ep-CoV2 forms a channel with K^+^ conductance. We further tested the Ep-CoV2 channel mutations V25F and N15A in the yeast complementation system as well. Both mutants appeared unable to rescue the growth phenotype under potassium-limiting conditions (Fig. 1H), which is in a general agreement with the aforementioned results from the suspended bilayer in that the N15A mutant is less active than the wt protein.

### The E-protein affects cytoplasmic calcium

During replication the structural membrane proteins of coronaviruses are synthesized into the ER membrane of infected host cells [38]. Since Ep-CoV2 has channel function in model membranes we reasoned that cells expressing this protein might exhibit an altered membrane conductance of the ER. This could be crucial in virus infected cells, since the ER is a major reservoir of important cell signaling factors like Ca^2+^ and H^+^.

In a first set of experiments, we addressed the question of whether Ep-CoV2 affects the intracellular concentration of free Ca^2+^ ([Ca^2+^]_in_). The viral protein was, therefore, expressed with or without a C-terminal GFP-tag in HEK293 cells. HEK293 cells were next co-transfected with cytosolic mRuby3 as a transfection marker with and without untagged Ep-CoV2. Representative images of control cells stained with the Ca^2+^ sensor Fluo4-AM (Fluo4) show the same Fluo4 signal in mRuby3 positive and negative cells indicating that the transfection marker is not affecting [Ca^2+^]_in_ (Fig. 2A,B) For cells co-transfected with Ep-CoV2 and mRuby3, we find that the Fluo4 signal is high in mRuby3-positive cells with respect to the mRuby3-negative controls (Fig. 2A,B). Given that the Fluo4 intensity approaches, but does not fully reach the values measured after adding the Ca^2+^-ionophore ionomycin (Fig. 2 A-C). Given that the Fluo4 reporter has a Kd value of 345 nM [Ca^2+^]_in_ [39] and is saturating at concentrations >1 µM, we can assume that the presence of Ep-CoV2 elevates [Ca^2+^] to a range of several hundred nM.

**Figure 2.**
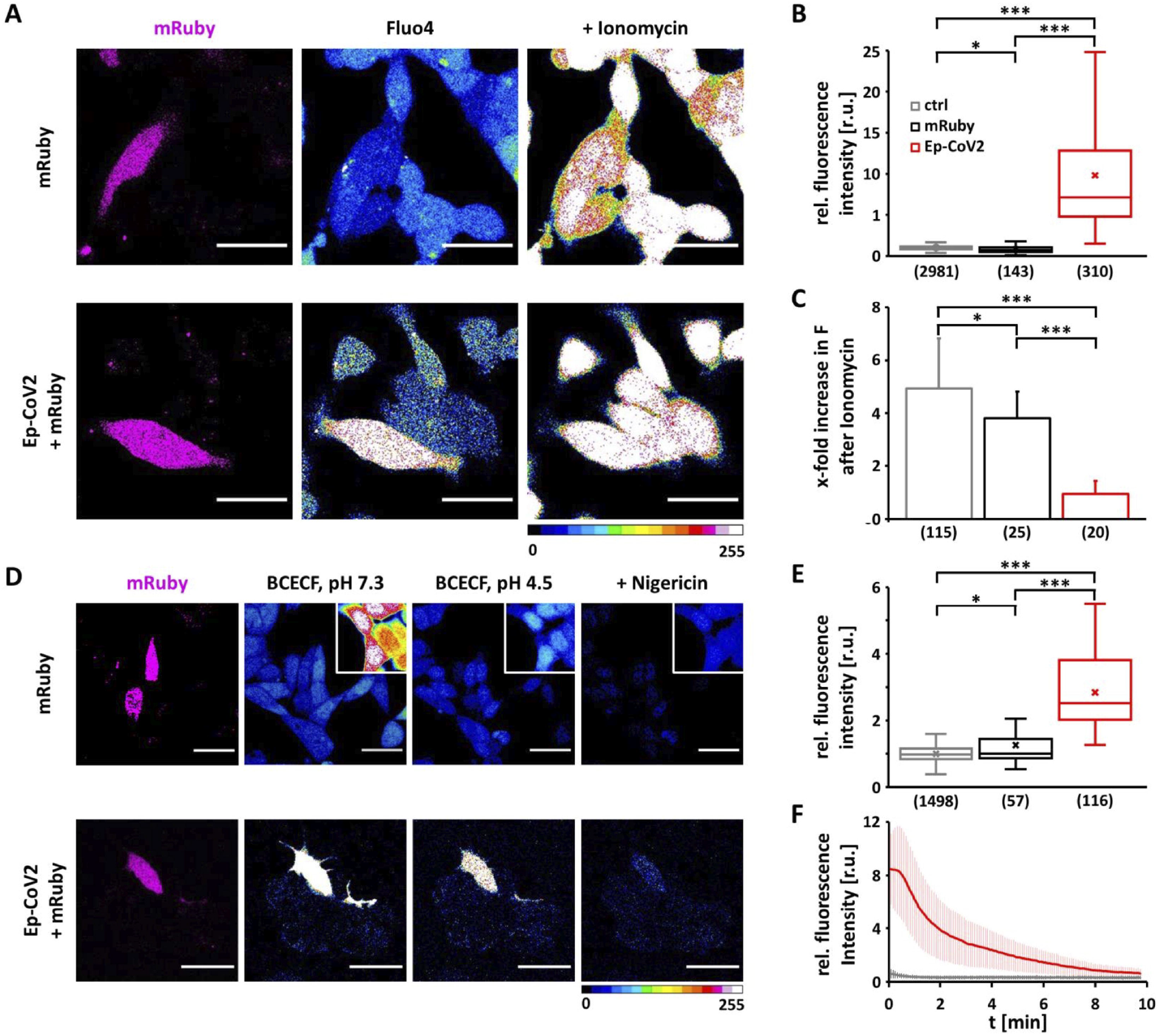
Expression of Ep-CoV2 in HEK293 cells increases [Ca^2+^]_in_ and pH_in_. **(A)** Representative fluorescence images of HEK293 cells transfected with either mRuby alone (upper row) or with mRuby + Ep-CoV2 (lower row). The signal of [Ca_2+_]_in_ reporter Fluo4 is globally low in control cells irrespectively of presence/absence of mRuby but high in cells transfected with mRuby + Ep-CoV2, lane two and three 16 color LUT. Addition of Ionomycin elevates [Ca_2+_]_in_ in all cells (lane three). Scale bar = 30 µm. **(B)** Boxplot of relative Fluo4 fluorescence intensity of HEK293 cells expressing mRuby alone (black) or Ep-CoV2 + mRuby (red) normalized to mean intensity of untransfected cells (ctrl, grey). Number of measured cells in brackets. **(C)** X-fold increase in Fluo4 fluorescence intensity after addition of Ionomycin. Bars represent arithmetic mean ± standard deviation; number of measured cells in brackets. **(D)** Representative fluorescence images of HEK293 cells transfected with either mRuby alone (first lane, magenta, upper row) or with mRuby + Ep-CoV2 (lower row). Fluorescence intensity of pH_in_ reporter BCECF at extracellular pH of 7.3 (second lane, 16 color LUT) or 4.5 (third lane, 16 color LUT) and after addition of nigericin (forth lane, 16 color LUT). BCECF signal is independent of presence/absence of mRuby in control cells but elevated in cells expressing mRuby + Ep-CoV2. While BCECF intensity decreases in control cells in acidic external buffer it remains high in cells expressing mRuby + Ep-CoV2. Nigericin causes in all cells a decrease in BCECF signal. Area framed in controls is shown at higher intensity to appreciate decay of BCECF signal. Scale bar = 30 µm. **(E)** Boxplot of relative BCECF-fluorescence-intensity of HEK293 cells expressing mRuby alone (black) or Ep-CoV2 + mRuby (red) normalized to mean intensity of untransfected cells (ctrl, grey); number of measured cells in brackets. **(F)** Time course of relative BCECF fluorescence intensity decay after addition of nigericin. Lines represent arithmetic means± standard deviations of 7 (Ep-CoV2, red) and 12 (ctrl, grey) cells. Boxes of the box plots in (B) and (E) represent the 25 ^th^ and 75^th^ percentile; the median is shown as horizontal line, the arithmetic mean as cross. The bars indicate the minimal and maximal value. Statistical significance in (B), (C) and (E) was determined with unpaired two-tailed student t-test assuming unequal variances (p < 0.001, ***; 0.001 < p < 0.01, **; 0.01 < p < 0.05, *; p > 0.05, n.s.).

We tested whether the increase in [Ca^2+^]_in_ is caused in an indirect manner by activation of endogenous plasma membrane Ca^2+^ channels. Experiments as in Fig. 2 were therefore repeated in the presence of conventional calcium-channel blockers Gd^3+^ (10 µM) or Bepridil (20µM). The presence of these blockers did not abolish the increased [Ca^2+^]_in_ in Ep-CoV2 expressing cells (Supplementary Fig. S2) implying that this rise in second messenger is not caused by an indirect activation of endogenous Ca^2+^ channels. The conclusion that Ca^2+^ influx across the plasma membrane is not the origin for the elevated [Ca^2+^]_in_ was confirmed by Mn^2+^-quenching experiments [40]. HEK293 cells transfected with Ep-CoV2 were therefore loaded with FURA2-AM and transferred into a buffer containing 2 mM Mn^2+^. This resulted in no apparent quenching of the FURA2 signal at 360 nm (Supplementary Fig. S.3) indicating that Mn^2+^ (as substitute for Ca^2+^) cannot enter the cells via the plasma membrane.

To test if the effect of Ep-CoV2 on [Ca^2+^]_in_ is a generic or cell-type-specific effect, experiments were also repeated in A549 cells, a model system for human alveolar cells which express the virus in infected humans [41]. Ep-CoV2 elicits also in these cells an increase in [Ca^2+^]_in_ (Fig. 5A,B) in these cells. Hence, expression of Ep-CoV2 causes an elevated [Ca^2+^]_in_ in a cell-type-independent manner, which is not mediated by endogenous plasma membrane Ca^2+^ channels. The results of these experiments are in a general agreement with recent reports showing that expression of Ep-CoV2 corrupts the Ca^2+^ homeostasis in mammalian cells and that this could be related to a release of the second messenger from internal stores [26]. Such a scenario could well explain the elevated [Ca^2+^]_in_, which we see in Ep-CoV2 expressing cells this study.

**Figure 3.**
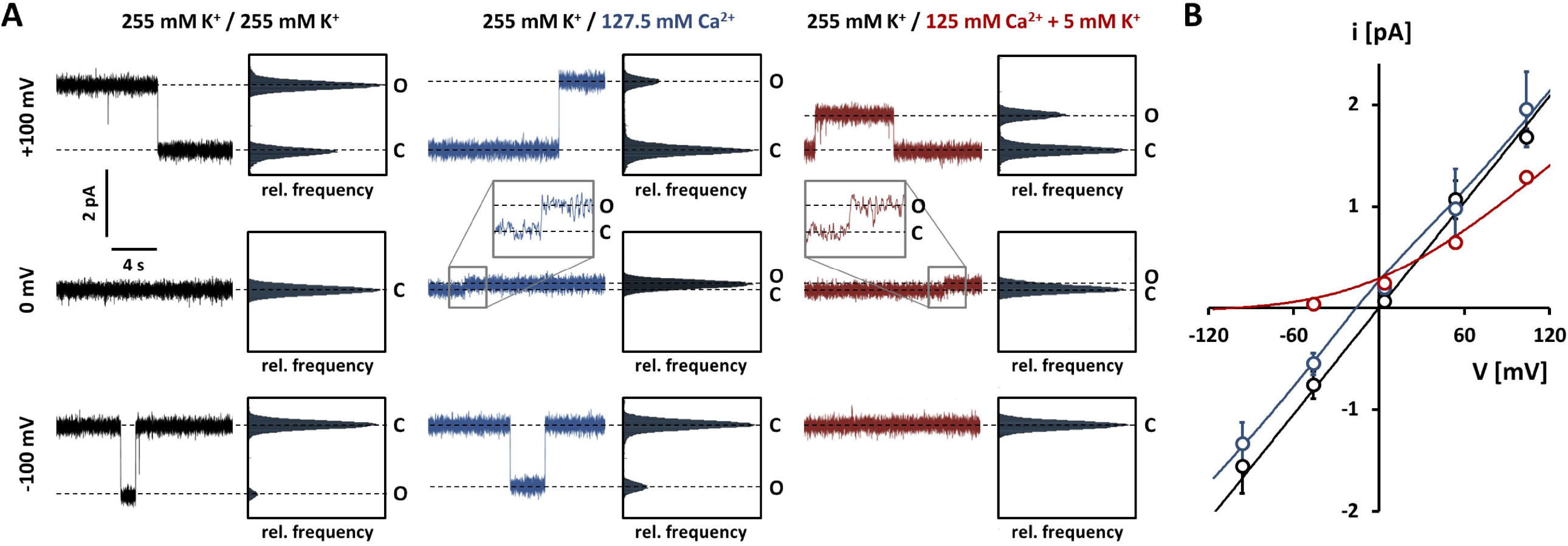
Ep-CoV2 shows Ca ^2+^ permeability and anomalous mole fraction effect in artificial lipid bilayers. **(A)** Representative Ep-CoV2 single channel current fluctuations in 1:1 DPhPC/DPhPS lipid bilayer at +100 mV, 0 mV and -100 mV. The solution in the *cis* chamber always contained 255 mM K^+^ (250 mM KCl, 1 mM EGTA, 10 mM HEPES, pH adjusted to 7.4 with ⍰5 mM KOH). The *trans* chamber was perfused with either 255 mM K^+^ (same as in *cis*) (left, black), 127.5 mM Ca^2+^ (125 mM CaCl_2_, 10 mM HEPES, pH adjusted to 7.4 with ⍰2.5 mM Ca(OH)_2_)(middle, blue), or 127.5 mM Ca^2+^ + 5 mM K_+_ (125 mM CaCl_2_, 10 mM HEPES, pH adjusted to 7.4 with ⍰5 mM KOH)(right, red). Corresponding amplitude histograms are depicted on the right of each trace. Dashed lines indicate the closed (C) or open channel (O) level. Current jumps measured under asymmetric conditions at 0 mV are shown enlarged in gray boxes above the corresponding traces. Current traces filtered at 100 Hz for visualization. **(B)** Unitary open-channel currents of Ep-CoV2 for the conditions shown in (A). Data points represent arithmetic mean ± standard deviation of at least three independent measurements. Solid lines were inserted for visualization purposes only and have no physical meaning.

**Figure 4.**
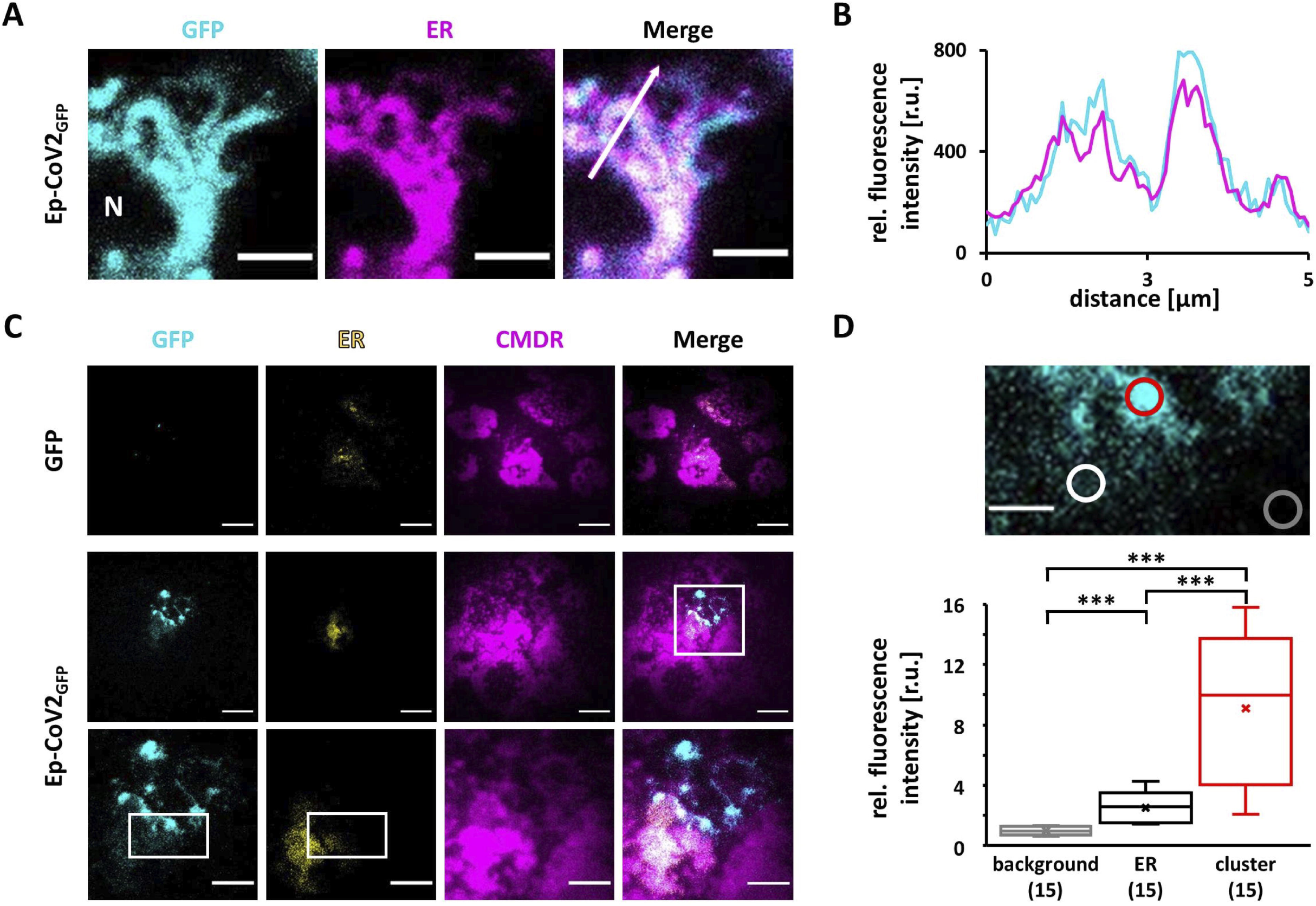
Ep-CoV2 is sorted into the ER of HEK293 cells. **(A)** Representative fluorescence images with zoom on ER in periphery of nucleus (N) in HEK293 cells transfected with GFP tagged Ep-CoC2 (Ep-CoV2_GFP_). GFP signal (cyan, left lane) and ER-tracker_™_ Blue-white DPX (magenta, middle lane) colocalize (right lane, merger). Scale bar = 3 µm. **(B)** Intensity profile along line in merger. Colocalization of free eGFP with ER signal (upper graph) or Ep-CoV2_GFP_ with ER signal (lower graph). **(B)** Plot profile along inserted arrow from merger in **A**. Colocalization of Ep-CoV2_GFP_ with ER signal. **(C)** Exemplary fluorescence TIRF microscopy images of isolated membrane patches of HEK293 cells expressing either eGFP (upper row) or Ep-CoV2_GFP_ (middle row). Area framed in merger is magnified in lower row. The GFP signal shows the localization of either eGFP or Ep-CoV2_GFP_ in isolated membrane patches (first lane, cyan). Remaining ER on the isolated patches (second lane) and isolated plasma membrane patch (third lane) are identified by ER-tracker_™_ Blue-white DPX (ER, yellow) and CellMask Deep Red (CMDR, magenta) fluorescence respectively. Mergers of lane 1, 2 and 3 show coincidence of Ep-CoV2_GFP_, ER and CMDR (fourth lane). Scale bars: 10 µm in upper and middle row or 3 µm in lower row. **(D)** Magnified area marked in GFP and ER channels with regions of high and low GFP intensity. Regions of interest (ROI, circles) positioned over region with high GFP/low ER (red) and low GFP/high ER intensity (white) as well as background with no GFP/ER fluorescence (grey). Relative fluorescence intensity of Ep-CoV2_GFP_ in background (grey), ER regions (black/white) and vesicular like clusters (red). Intensity was normalized to mean background intensity. Boxes in box plots in (D) represent the 25^th^ and 75^th^ percentile; the median is shown as horizontal line, the arithmetic mean as cross. Bars indicate the minimal and maximal value. Scale bar = 2 µm. Statistical significance in (D) was determined with unpaired two-tailed student t-test assuming unequal variances (p < 0.001, ***; 0.001 < p < 0.01, **; 0.01 < p < 0.05,*; p > 0.05, n.s.).

**Figure 5.**
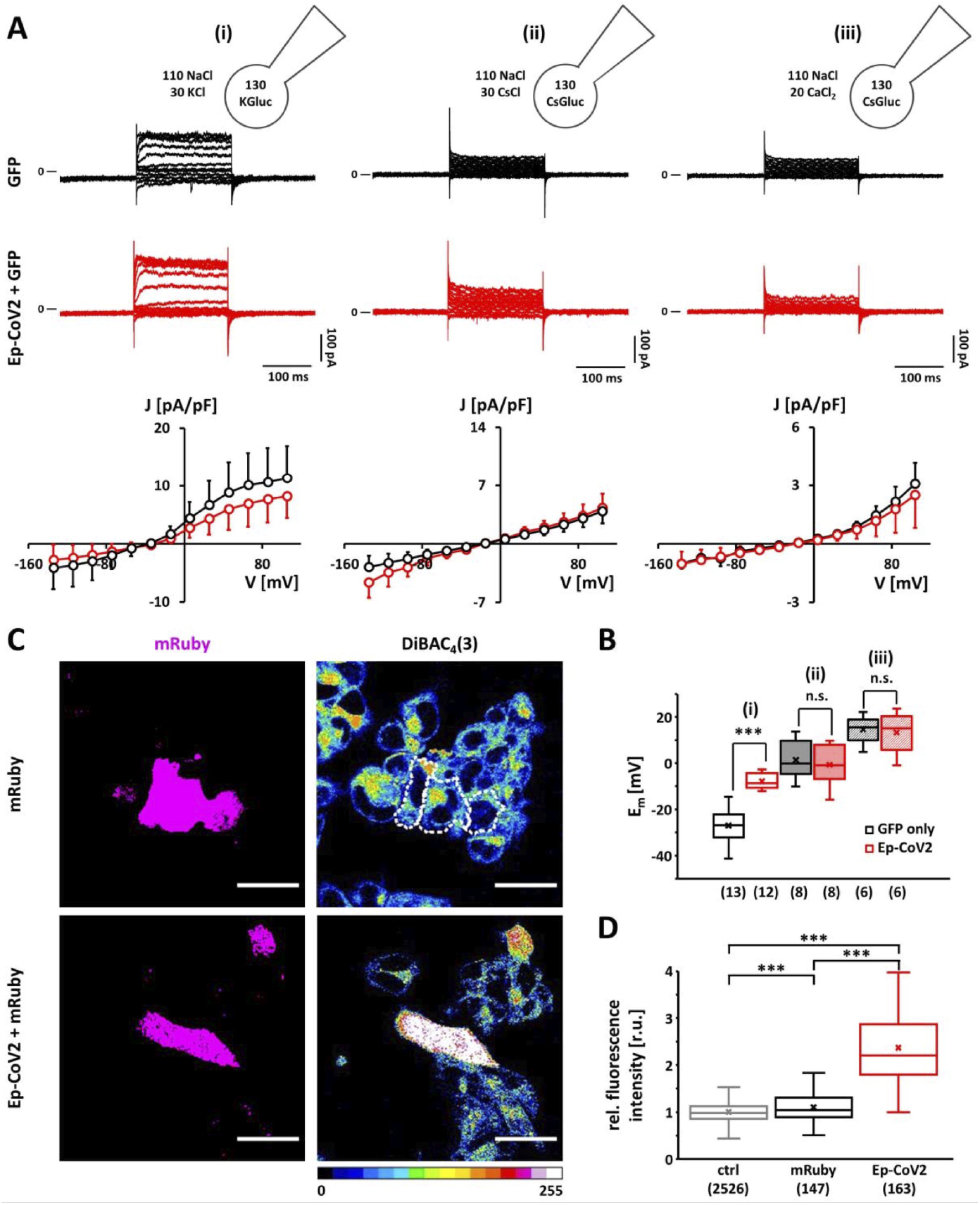
Expression of Ep-CoV2 has no appreciable effect on whole-cell currents but depolarizes the plasma membrane of HEK293 cells. **(A)** Whole-cell voltage-clamp recordings of HEK293 cells expressing GFP (negative control, upper row in black) or Ep-CoV2 and GFP (middle row in red). Whole-cell currents elicited by voltage steps between -135 mV and +105 mV from holding potential of -55 mV. Current density (J) / voltage relations (lower row) represent mean current densities at respective clamp voltages. Recordings were performed using standard internal and external solutions (i), solutions containing cesium instead of potassium (ii), and solutions containing cesium and high external CaCl_2_ (iii). For more details see materials and methods. **(B)** Mean membrane potentials (E_M_) of HEK293 cells expressing GFP (black) or Ep-CoV2 + GFP (red) measured by current-clamp in whole-cell configuration in solutions (i) - (iii) as in (A). Number of measured cells in brackets. **(C)** Representative fluorescence images of HEK293 cells expressing either mRuby alone as negative control (upper row) or Ep-CoV2 + mRuby (lower row). mRuby signal acts as transfection control (first lane, magenta). Cells were loaded with 10 µM DiBAC_4_(3) to monitor membrane potential (lane two, 16 color LUT). Contours of mRuby positive cells in DiBAC_4_(3) channel in upper row are highlighted with white dotted line. Scale bar = 30 µm. **(D)** Relative DiBAC_4_(3) fluorescence intensity of HEK293 cells expressing mRuby alone (black) or Ep-CoV2 + mRuby (red) normalized to the mean intensity of untransfected cells (ctrl, grey); number of measured cells in brackets. The boxes of the box plots in (B) and (D) represent the 25^th^ and 75^th^ percentile; the median is shown as horizontal line, the arithmetic mean as cross. The bars indicate the minimal and maximal value. Statistical significance in (B) and (D) was determined with unpaired two-tailed student t-test assuming unequal variances (p < 0.001,***; p > 0.05, n.s.).

### E-protein in model membranes shows an anomalous mole fraction effect

To further test if this increase in [Ca^2+^]_in_ is directly mediated by the channel activity of Ep-CoV2 we examined the Ca^2+^ conductance in planar lipid bilayers. Channel activity was first measured in a symmetrical 250 mM KCl solution before the buffer on the *trans*-side was exchanged with a K^+^-free 125 mM CaCl_2_ solution (Fig.3A left panel). In this condition we measured positive and negative channel fluctuations (Fig. 3A,B). This suggests that the channel conducts K^+^ outward and Ca^2+^ inward currents. When the solution on the *trans*-side, in the same recording, contained 125 mM CaCl_2_ and 5 mM K^+^ we only observed K^+^ outward current with no appreciable inward current (Fig. 3A middle panel, Fig3B). The results of these experiments suggest that the channels show an anomalous mole fraction effect (AMFE), a phenomenon that was already reported for a higher unitary conductance of Ep-CoV2 [16]. The results of these experiments suggest that Ep-CoV2 conducts Ca^2+^ and K^+^ in pure solutions. A mixture of the two ions is not additive but they cause a mutual inhibition of unitary conductance. These data indicate that Ep-CoV2 can, under specific constraints, conduct Ca^2+^; however, we cannot determine whether Ep-CoV2 is solely responsible for the increase in [Ca^2+^]_in_ observed under physiological conditions. Instead, our data suggests an indirect effect of Ep-CoV2 on the regulation of endogenous Ca^2+^ homeostasis.

### The E-protein has an impact on cellular pH homeostasis

To further determine the effect of Ep-CoV2 on cellular ion homeostasis, we sought to identify any possible effects on cellular pH (pH_in_); to this end we performed experiments using the pH sensitive dye BCECF-AM. Confocal images of control cells expressing only mRuby3 exhibit a homogeneous signal in all cells irrespectively of reporter expression (Fig. 2D,E). In cells co-transfected with mRuby3 and Ep-CoV2, we found a slight increase in the fluorescent signal throughout the cell suggesting that Ep-CoV2 causes alkalization of the cytosol. To further test the significance of this effect the external buffer was exchanged from a neutral (pH 7.3) to an acidic (pH 4.5) medium (Fig. 2D). The consequent acidification of the cytosol resulted in mRuby3 negative cells in a slow decrease in the fluorescent signal. In contrast, mRuby3 positive cells, which were co-transfected with Ep-CoV2, maintained a high fluorescent signal (Fig. 2D,E). The results of these experiments suggest that Ep-CoV2 is not generating a proton conductance in the plasma membrane, but it is instead supporting cellular systems, which buffer the cytosolic pH close to its resting value. In both Ep-CoV2 transfected and un-transfected cells the intracellular pH decreased to a common level when the H^+^ conductance of the plasma membrane was increased by nigericin; while this acidification was rapid in control cells it progressed much slower in Ep-CoV2 positive cells (Fig. 2D). Hence the pH-buffering effect of Ep-CoV2 can be eventually overridden by a massive influx of H^+^ across the plasma membrane.

### Ep-CoV2 is located at the plasma membrane and affects the membrane potential

There are controversial reports on whether the E-protein from SARS-CoV1 and SARS-CoV2 can also form ion channels in the plasma membrane of cells. While some studies exclude an Ep-CoV1 generated conductance in the plasma membrane [25,26] others find elevated currents after expressing the protein in the same system [19,24]. A recent publication suggests that the presence or absence of the Ep-CoV2 in the plasma membrane is a matter of protein sorting. Only after addition of an appropriate sorting signal was Ep-CoV2 trafficking to the plasma membrane in the case of *Xenopus* oocytes and mammalian cells generating a cation selective conductance [25].

Since the aforementioned yeast complementation assays suggested that Ep-CoV2is present in the plasma membrane (Fig. 1), we attempted to visualize the protein in the plasma membrane of HEK293 cells. When GFP-tagged Ep-CoV2 (Ep-CoV2_GFP_) was expressed in HEK293 cells, the fluorescence associated with the protein co-localized, like in other studies [25,26], well with the fluorescent marker ER-Tracker™ which stains specifically the endoplasmic reticulum (Fig. 4A,B). To address the question of a plasma membrane localization for the Ep-CoV2, we used high-resolution microscopy of membrane proteins in the plasma membrane [42]. Small plasma membrane patches of HEK293 cells expressing Ep-CoV2_GFP_ were isolated by deroofing of the cell body [43] and imaged with TIRF microscopy. Fig. 4C shows representative images of such plasma membrane (PM) patches transfected with either GFP alone or with Ep_GFP_-CoV2. The isolated PM patch in the evanescent field can be identified from the fluorescence of the PM-specific dye CellMask™ Deep Red (CMDR, magenta). From the uneven intensity of this dye we must conclude that the membrane patch is not making a flat foot over the entire contact with the cover slip. In some cases, like in the example in Fig. 4C, small areas with a fluorescent signal from the ER-tracker™ (yellow) are visible, suggesting some remaining cortical ER attached to the patches.

In cells expressing only soluble GFP, we never detected any GFP fluorescence in the evanescent field of the isolated membrane patch or the associated ER. Cells expressing Ep-CoV2_GFP_, on the other hand, frequently show a GFP signal in the plane of the PM; this signal is composed of intense spots and, in the case of remaining cortical ER, of a diffuse low-intensity background (Fig. 4C,D). In the overlay, the diffuse low-background GFP signal colocalizes with the ER marker, suggesting that the protein is present in the cortical ER. The intense fluorescent GFP spots, on the other hand, colocalize exclusively with the PM marker (Fig. 4C). These spots of colocalization are never in the foot area where the PM touches the cover slip but higher up in the evanescent field. At this point, we cannot distinguish if these are specific domains of the membrane or if they are vesicular structures (e.g. exosomes) which were secreted from the cell. The same patterns of fluorescent signal distributions were detected in six other cells expressing Ep-CoV2_GFP_. Typically, the GFP signals were slightly but significantly elevated above the background in areas, which revealed a signal from the ER (Fig. 4C,D). In ER-free areas, the GFP signal showed a spot-like distribution with an intensity well above that associated with the ER (Fig. 4C,D). The results of these experiments confirm that Ep-CoV2_GFP_ is present in the ER and reaches distinct areas of the plasma membrane, where it occurs at high concentration in a patch-like manner.

With the indication of a potential PM localization of Ep-CoV2_GFP_, we tested its channel conductance by whole-cell patch-clamp measurements in HEK293 cells. Comparative inspection of representative current responses with corresponding mean steady-state I/V relations from cells transfected with GFP or co-transfected with GFP and Ep-CoV2 shows no obvious indication for a conductance of Ep-CoV2 at the plasma membrane (Fig. 5A). The same is true for cells transfected with the GFP-tagged version of Ep-CoV2 (Ep-CoV2_GFP_). The observed similarity in the I/V characteristics between control cells and cells expressing Ep-CoV2 is present not only for measurements in physiological solutions containing K^+^ and Na^+^ but also in solutions with Cs^+^ or high external Ca^2+^ (Fig. 5A). Hence, in line with data from others [26], Ep-CoV2 does not generate a measurable cation conductance in the plasma membrane of HEK293 cells. Either the protein is not present as a functional channel in the plasma membrane or the additional Ep-CoV2 conductance is smaller than the scatter of current densities between individual cells.

While the data show no evidence for an Ep-CoV2-mediated conductance at the plasma membrane, they do indicate that the viral protein causes a depolarization of the membrane potential in a K^+^ containing solution. Close inspection of the I/V characteristic from cells expressing Ep-CoV2 shows that the reversal voltage is depolarized and shifted by about 9 mV to a membrane potential more positive than the controls. In complementary experiments, we measured the resting membrane potentials (E_M_) of cells ± Ep-CoV2 in current clamp recordings. The data in Fig. 5B confirm that expression of Ep-CoV2 causes a depolarization of the membrane potential (E_M_) of 19 mV. This effect was only evident when Ep-CoV2 was expressed without a GFP tag and in recordings with K^+^ in the internal and external solutions. Substitution of K^+^ with Cs^+^ resulted in a strongly depolarized membrane potential in both Ep-CoV2-transfected (E_M_ = -0.7 ± 8.7 mV) and control cells (E_M_ = +1.3 ± 8.2 mV) (Fig. 5B). Further elevation of the external Ca^2+^ concentration from 2 to 20 mM did not result in a difference in membrane potentials between Ep-CoV2-transfected (E_M_ = +9.2 ± 13.8 mV) or control cells (E_M_ = +13.4 ± 8.2 mV), providing additional evidence that the positive shift of the membrane potential in Ep-CoV2-expressing cells is mediated by inhibition or elimination of an endogenous K^+^ conductance or a K^+^-dependent electrogenic transport process.

The same measurements were repeated in A549 cells revealing a much larger variability of currents among control cells (Fig. 6 E,F). Typically, these cells exhibit different degrees of inward currents as well as outward currents with different types of kinetics. The latter are either dominated by an inactivating and/or non-inactivating outward rectifier. As much as the control cells show different types of currents, the membrane potentials of these cells are also more variable, including cells with strongly hyperpolarized and depolarized voltages (Fig. 6G). The mean E_M_ from 24 different control cells was -13.6± 6.3 mV.

**Figure 6.**
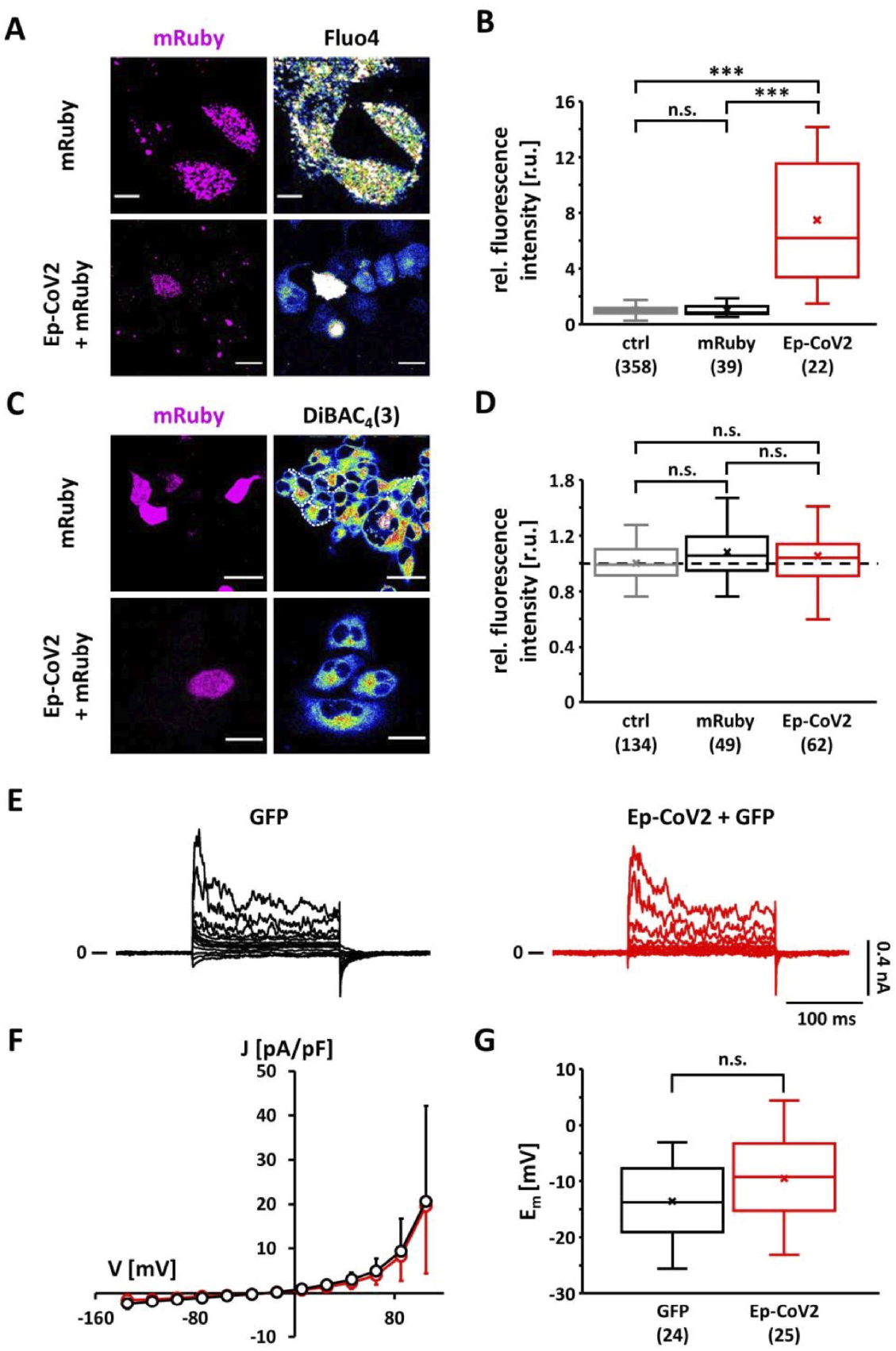
Expression of Ep-CoV2 increases [Ca^2+^]_in_ of A549 cells but has no appreciable effect on electrical properties of the plasma membrane. **(A+C)** Representative fluorescence images of A549 cells expressing either mRuby alone (negative control; upper row) or mRuby plus Ep-CoV2 (lower row). In both cases mRuby signal acts as a transfection control (first lane, magenta). Cells were loaded with 1 µM Fluo4-AM to indicate the [Ca^2+^]_in_ **(A)** or 10 µM DiBAC_4_(3) to monitor their membrane potential **(C)** (second lane, 16 color LUT). Fluo4 intensity is insensitive to absence/presence of mRuby but elevated in cells co-transfected with mRuby + Ep-CoV2. DiBAC_4_(3) fluorescence is independent on presence/absence of mRuby or mRuby + Ep-CoV2. Contours of mRuby positive cells in DiBAC_4_(3) channel in upper row are highlighted with white dotted line. **(B+D)** Relative Fluo4 **(B)** or DiBAC_4_(3) **(D)** fluorescence intensity of HEK293 cells expressing mRuby alone (black) or Ep-CoV2 + mRuby (red) normalized to mean intensity of untransfected cells (ctrl, grey); number of measured cells in brackets. **(E)** Representative whole-cell voltage-clamp recordings of A549 cells expressing GFP (black) or Ep-CoV2 and GFP (red) using standard internal and external solutions. Currents were elicited by voltage steps between -135 mV and +105 mV from holding potential of -55 mV. Corresponding mean current density/voltage relations at respective clamp voltages are shown in **(F)** (GFP, black, N = 24; Ep-CoV2+GFP, red, N = 25). **(G)** Mean membrane potentials (E_m_) of A549 cells expressing GFP or Ep-CoV2 and GFP measured by current-clamp in whole-cell configuration in standard solutions; number of measured cells in brackets. Boxes of the box plots in (B), (D), and (G) represent the 25^th^ and 75^th^ percentile; the median is shown as horizontal line, the arithmetic mean as cross. The bars indicate the minimal and maximal value. Statistical significance in (B), (D) and (G) was determined with unpaired two-tailed student t-test assuming unequal variances (p < 0.001,***; p > 0.05, n.s.). Scale bars represent a distance of 10 µm (A, upper row), 20 µm (A, lower row + C, lower row) and 30 µm (C, upper row).

Electrophysiological recordings of cells expressing Ep-CoV2 show the same heterogeneity of membrane currents as the controls (Fig. 6E,F). The mean I/V characteristics of Ep-CoV-expressing cells match the heterogeneity of currents observed in the controls. In agreement with our results using HEK cells, this suggests that Ep-CoV2 is not generating an appreciable ion conductance at the plasma membrane of A549 cells. The E_M_ of 25 Ep-CoV2-expressing cells is -9.5 ±7.9 mV showing a non-significant (p =0.06) depolarization compared to the control cells (Fig. 6G).

The effect of Ep-CoV2 on the membrane voltage of the two cell lines was confirmed by fluorescent measurements in cells with and without Ep-CoV2. Quantitative image analysis show that expression of mRuby3 alone has no impact on the fluorescence intensity of the voltage sensitive dye DiBAC_4_(3) in both cell types (Fig. 5C,D; 6C,D). However, when HEK293 cells were co-transfected with mRuby3 and Ep-CoV2, the mRuby3-positive cells exhibited a significant increase in DiBAC_4_(3) fluorescence (Fig. 5C,D). This decrease in fluorescence indicates a depolarized membrane voltage. We confirmed these findings using the fluorescent voltage reporter FluoVolt. The results (Supplementary Fig. S4) are overall in good agreement with the electrical recordings, showing a depolarized E_M_ in Ep-CoV2-expressing cells. In contrast, A549 cells show no appreciable difference in DiBAC_4_(3) fluorescence between cells with or without Ep-CoV2, which is also in agreement with the electrophysiological recordings.

Taken together the data provide evidence that Ep-CoV2 affects the membrane potential of mammalian cells in a cell-type-specific manner. We are unable to detect any significant change of membrane currents above the measurement noise level in Ep-CoV2-expressing cells that would explain this shift of the membrane potential. These data indicate that only a small number of channels is reaching the plasma membrane where they generate a conductance that is in the range of the variability of current densities between individual cells. This interpretation would be consistent with the very low unitary conductance seen in bilayer recordings (Fig 1) and the need of a targeting signal for efficient trafficking of Ep-CoV2 to the plasma membrane [25]. The explanation would also agree with the finding that expression of Ep-CoV2 in yeast is sufficient for rescuing yeast growth (Fig. 1F-H). An alternative interpretation, which cannot be excluded at this point, is that the viral protein has no direct channel activity in the plasma membrane but modulates a transporter, which is strongly contributing to the membrane voltage in HEK293 but not in A549 cells. While this interpretation is compatible with the data in mammalian cells, it is not in agreement with the finding that Ep-CoV2 is able to rescue K^+^ uptake deficient yeast.

### Mutants alter the impact of Ep-CoV2 on cellular parameters

After finding that the expression of Ep-CoV2 in HEK293 cells modulates [Ca^2+^]_in_, pH_in_ and E_M_, three distinct cellular functions which are all related to cellular signaling cascades, we asked the question whether these parameters are correlated and whether they are directly or indirectly related to a channel function of the viral protein. To answer this question, we repeated ([Ca^2+^]_in_, pH_in_ and E_M_ measurements with mutant constructs of Ep-CoV2. Based on functional considerations deducted from homology modeling and from previously reported experimental studies, we assume that a C- and N-terminal truncation of (Ep-CoV2_™_) (Fig. 1A) should eliminate interactions with endogenous proteins [44] while not compromising channel function [14]. However, following observations in Ep-CoV1, point mutations N15A and V25F in the transmembrane domain of the proteins N15A and V25F should eliminate or impair channel function [14, 22] by corrupting oligomerization of Ep-CoV2 [3]. A recent study on full-length Ep-CoV2, however, reports that both of these mutations affect the structure of the protein to differing extents but not its ability to oligomerize [11]. In order to access the impact of these mutations on channel function, we repeated the aforementioned yeast complementation assay with the respective Ep-CoV2 mutants. The data show that none of the mutants was able to rescue yeast growth (Fig. 1H). This implies that all the mutations including the truncation of cytosolic/extracellular domains, must have a strong impact on channel function at least in the yeast system.

Functional testing of these mutants in HEK293 cells shows that the rise in [Ca^2+^]_in_ is reduced, but not eliminated, by expressing either the truncated Ep-CoV2 protein or the two mutants (Fig. 7A). It is worth noting that the Ep-CoV2-V25F mutant, which should have the most severe impact on ion channel function [3], does not behave differently than the Ep-CoV-N15A mutant and the C- and N-terminal-truncated Ep-CoV2_™_. A similar picture emerges from the pH recordings; here, neither the truncated protein nor the mutants have a strong impact on the amplitude of the pH excursion in cells which express Ep-CoV2 (Fig. 7B). Collectively the data show that mutations, which are thought to alter the structural properties of Ep-CoV, have an impact on its effect on cellular functions, which is not correlated with their impact on protein oligomerization and channel function [3]. This implies that the rise in [Ca^2+^]_in_ and pH either is not caused primarily by a channel function of Ep-CoV2 or that the impact of the mutations from *in vitro* experiments cannot be directly extrapolated to functional data in cells. It is possible that mutations N15A and V25F impair (but do not eliminate) oligomerization and, consequently, channel function. The latter interpretation is agreement with our findings that the N15A mutant was still able to generate some channel functions in artificial membranes (Fig. S1C) and that both mutants exhibit the same oligomerization as the full-length Ep-CoV2 wt protein [11].

**Figure 7.**
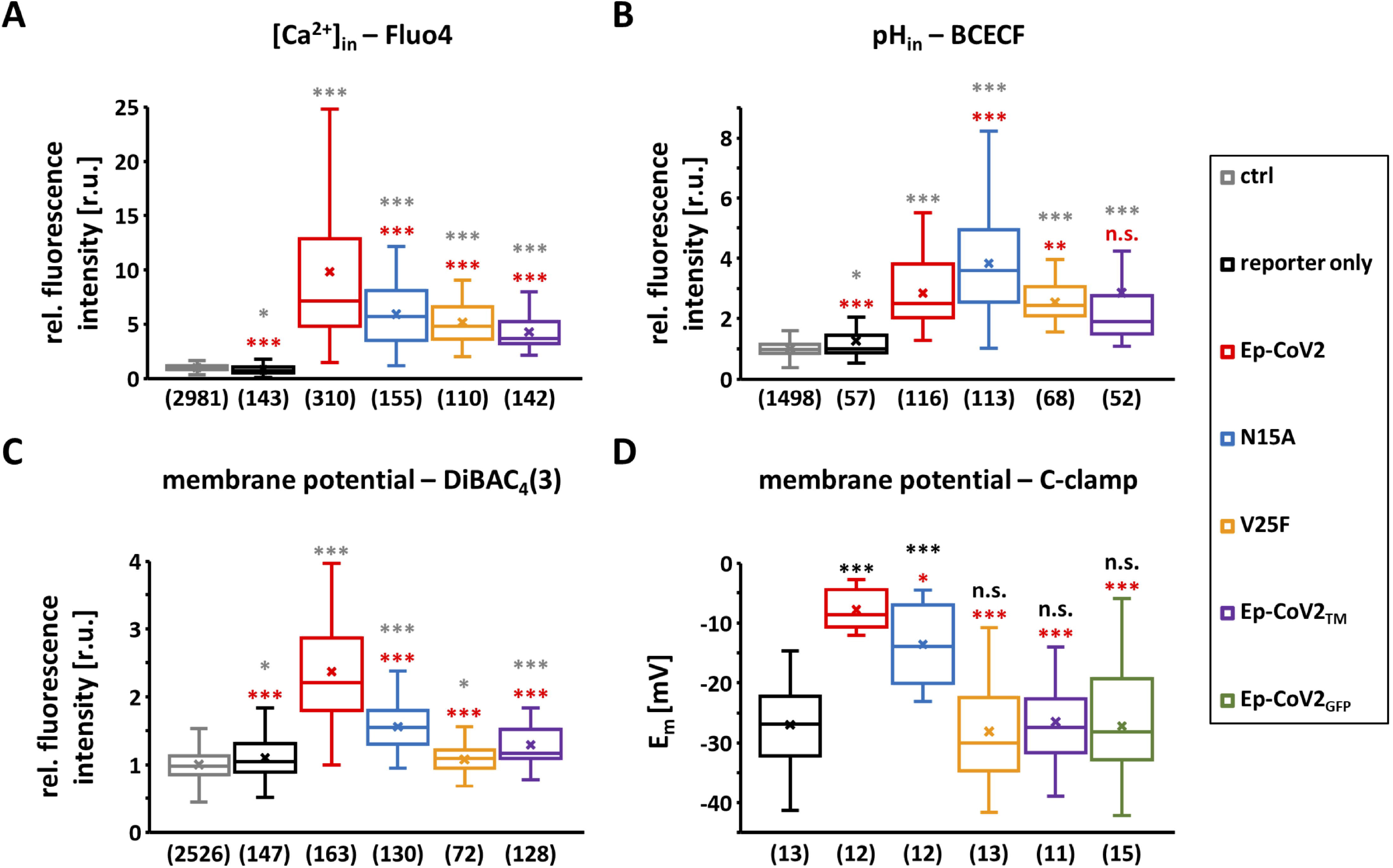
Effects of Ep-CoV2 and its mutants on [Ca^2+^]_in_, pH_in_ and membrane potential. **(A-C)** Box plot of relative fluorescent intensities from sensors for **(A)** Ca^2+^(Fluo4), **(B)** pH (BCECF) and **(C)** membrane voltage (DiBAC4(3)) in HEK293 cells expressing mRuby alone (black) or mRuby + Ep-CoV2 variants. Red (wt), blue (N15A), yellow (V25F) and purple (Ep-CoV2TM). Data normalized to mean intensity of un-transfected control-HEK293 cells (grey); number of measured cells in brackets. **(D)** Mean membrane potentials (E_M_) of HEK293 cells expressing eGFP (black), eGFP + Ep-CoV2 variants. Red (wt), blue (N15A), yellow (V25F) and purple (Ep-CoV2_™_) or green Ep-CoV2_GFP_. E_M_ were measured by current-clamp in whole-cell configuration; number of measured cells in brackets. Boxes of box plots in A-D represent 25^th^ and 75^th^ percentiles; median is shown as horizontal line, arithmetic mean as cross. Bars indicate minimal and maximal value. Statistical significance was determined with unpaired two-tailed student t-test assuming unequal variances (p < 0.001, ***; 0.001 < p < 0.01, **; 0.01 < p < 0.05,*; p > 0.05, n.s.).

The impact of the mutants on E_M_ were again examined by fluorescent measurements and by patch-clamp recordings in HEK293 cells. The general picture which emerges from both assays suggests that the mutant proteins either reduce (EP-CoV2-N15A) or eliminate (EP-CoV2-V25F and Ep-CoV2_™_) the depolarizing effect of Ep-CoV2 (Fig. 7C,D). The mutants affecting the oligomerization of the protein (EP-CoV2-V25F) or its interaction with other protein partners (Ep-CoV2_™_) fully eliminate the depolarizing effect of Ep-CoV2 on the membrane voltage (Fig. 7C,D). Again the effects of the mutations on membrane depolarization are not in line with their expected impact on channel function, meaning that either biochemical data cannot be extrapolated in channel function or that the depolarization is not the result of a channel function. The data would also support a scenario in which Ep-CoV2 interacts with either a pump or a channel; the resulting modulation of the latter is then causing the membrane depolarization. It is worth noting that such a scenario could also explain the negative effect of the GFP tag on Ep-CoV2-induced membrane depolarization because it could sterically hinder the proposed interaction with other proteins.

From the differential impact of the mutants on either [Ca^2+^]_in_ and pH_in_, compared to their effects on membrane potential on the other side (Fig. 7A-D), we can further conclude that the membrane depolarization in Ep-CoV2 expressing cells is neither a secondary result of altered Ca^2+^ nor a result of pH conditions. The apparent requirement of this effect on the cytosolic terminus (Fig. 7C,D) instead suggests that interactions of this domain with cellular factors are essential for this effect. Such interactions could also be disturbed in a case where the oligomerization of the protein is corrupted. It is possible that in this case the extracellular/cytosolic termini are misfolded preventing an interaction with partner proteins.

## DISCUSSION

Host cells infected with coronaviruses produce abundant amounts of E-protein, of which only a small part is incorporated into the nascent virus particle as structural membrane protein; the majority of the proteins remain in host cell membranes. The data presented here shed light on the potential impacts of the latter protein fraction on functional properties of infected cells. A key concept of cell-signaling is that crucial regulatory factors are under a homeostatic control. Any deviation from a narrow set-point affects complex signaling cascades, which modulate a large spectrum of functions ranging from acute reactions to long lasting alteration in cell development and differentiation. This concept has been extensively studied for Ca^2+^ and pH [45, 46]. Here we find that expression of the E-protein affects three of the main homeostatic control systems of mammalian cells -- the plasma membrane voltage, [Ca^2+^]_in_ and pH_in_. In this way, CoVs follow a strategy that is used by many other viruses in that they commandeer universal signaling pathways in their host cells to their own benefits. The three signaling parameters, which are modulated by the E-protein, are so universal in cells that any of the pathophysiological reactions in infected cells and tissues can potentially be associated with signaling cascades, which are triggered in response to E-protein activity. The present data on an E-protein-mediated increase in Ca^2+^ are interesting in the context of a recent report on the Porcine deltacoronavirus (PDCoV). It was found that this virus, which is related to SARS-CoV1 and SARS-CoV2, causes an increase in [Ca^2+^]_in_ in host cells [47]. Further experiments revealed that pharmacological manipulations, which reduced the rise in [Ca^2+^]_in_, were able to inhibit distinct steps of PDCoV replication. In the context of these findings, it is tempting to speculate that the effects of Ca^2+^-channel blockers [48,49] and Ca^2+^ chelators [50] on reducing mortality from Covid-19 could be directly or indirectly related to the function of the E-protein in elevating [Ca^2+^]_in_.

The data presented here cannot answer the question on whether the effects of the E-protein on the signaling systems are solely mediated by its channel function or in indirect response to such signal cascades. Our data confirm reports by others that the E-protein is indeed generating channel activity in artificial membranes. In our experimental approach, the protein is translated *in vitro* in the presence of an artificial target membrane, the lipid nanodisc, which resembles the native situation in which the protein is produced at the ER membrane. Under our experimental conditions the protein generates a highly reproducible channel activity with a small unitary conductance of < 20 pS and remarkable long open and closed dwell times with seconds-long duration. Large conductances, which were reported by others, were not observed. One possible explanation for this discrepancy with other studies, is that the delimited size of the nanodiscs may function as a size exclusion filter, which would only allow the formation of a defined oligomeric state. In other conditions, the protein may also form higher order oligomers, which may generate larger unitary conductances.

Our data suggest a plasma membrane localization of the E-protein-generated channel. The most convincing evidence comes from the yeast-complementation assay in which the expression of the E-protein is able to rescue growth of a K^+^-uptake-deficient mutant. This is best understood in a scenario in which the cation permeable E-protein mediates a K^+^ conductance in the plasma membrane, which is sufficient for providing K^+^ influx in a selective medium with low-potassium concentration. The interpretation of the data in mammalian cells is less straightforward but can also be explained in the same manner. High-resolution imaging data suggest that GFP-tagged E-protein is present in the plasma membrane in distinct clusters. The limits of the resolution, however, do not allow any detailed identification of these structures. The peculiar positioning does not even exclude that these structures are of vesicular nature inside or outside of the membrane. Electrophysiological recordings and fluorescent imaging show that expression of the E-protein causes a distinct depolarization of the membrane voltage in HEK293 cells. This effect cannot be directly correlated to an increase in an E-protein-mediated channel function in the plasma membrane.

There are at least two possible explanations for this depolarization; it has been shown that the E-protein is only trafficked efficiently to the plasma membrane after adding a targeting sequence [25]. This does not exclude the possibility that this system is leaky, that is, that a small number of channels is reaching the plasma membrane. Since the viral proteins and the mammalian cells did not undergo co-evolution, it is most likely that such a sorting system is not perfect. A small amount of E-protein-formed channels may be sufficient to depolarize the membrane. At the same time, a small number of channels with small unitary conductances can also remain undetected when comparing different populations of cells, e.g. control cells and cells expressing the E-protein.

While this scenario is plausible the data cannot exclude an alternative explanation in which the effects on the membrane voltage are not an immediate cause of a channel function but an indirect effect of the E-protein. It has been reported that the transmembrane domain of the E-protein by itself is sufficient to generate channel activity [14]. If this is also true in a cellular context, the present data argue against a causal relation between channel activity and membrane depolarization. Notably, the truncated mutant of the E-protein, Ep-CoV2_™_, which should still have channel activity, is not able to evoke a membrane depolarization (Fig. 7C,D). In the light of these data, we cannot exclude the possibility that the E-protein interacts as a monomer or oligomer with other transport proteins in the plasma membrane, which contribute to the membrane potential of these cells. One potential candidate for such an interaction could be the K^+^/Na^+^ ATPase. It has been reported that the E-protein from SARS-CoV1 interacts with the Na^+^/K^+^ ATPase [51], an active pump responsible for the hyperpolarization of the free running membrane voltage in mammalian cells. Support for such an alternative explanation based on indirect effects of Ep-CoV2 with transport proteins comes from the fact that truncation of the C-terminus, a domain known to promiscuously interact with other proteins via the well-studied PDZ-binding motif (PBM) [52] (Fig. 1A), is effective in eliminating the impact of the E-protein on membrane voltage.

The present data also suggest a causal relationship between channel function of the E-protein and its modulation of [Ca^2+^]_in_ and pH_in_. The electrophysiological recordings imply that the channel has some Ca^2+^-permeability which is suppressed by K^+^ in the medium, suggesting an anomalous mole fraction behavior of the channel, a feature that has already been reported for the E-protein channel from SARS-CoV1 [16]. However, the unitary channel conductance in the latter study is very different from the one that we measure in our study; it is not clear if both studies look at the same mechanism. Because of an apparent complex interplay between K^+^ and Ca^2+^ on the permeability of the channel, we cannot easily predict from the *in vitro* measurements whether the Ep-CoV2 channel conducts Ca^2+^ in the ER membrane under physiological conditions. However, because of the very low concentration of Ca^2+^ in the cytosol, a small and steady Ca^2+^ leak in the ER membrane could already be sufficient to cause the observed increase of [Ca^2+^]_in_. Further systematic studies of the functional properties of the Ep-CoV2-generated channel are needed to understand its full impact in the complex cell environment.

As in the case of membrane depolarization, the experiments with E-protein mutants do not explain why elevated [Ca^2+^]_in_ and pH_in_ are observed. These effects are reduced in the case of mutations which are know to inhibit channel function. These same effects are also observed, however, with truncated proteins, which were thought to preserve channel function [3]. In the context of these data, we conclude that these effects on [Ca^2+^]_in_ and pH_in_ are sensitive to E-protein structural mutations, but additional studies are needed to answer the question of whether the effect of the E-protein is caused by a channel function or is due to an indirect effect of the monomeric or oligomeric form on cellular proteins.

Even though the mode of action of the E-protein in cells is not fully understood, the present data underpin that the viral protein modulates cellular functions in multiple ways. With such a prominent role in infected cells and with the known importance of the E-protein in the pathogenicity of the virus this protein presents a key target for developing virostatic drugs. This choice as a target is further supported by the fact that the E-protein exhibits a much lower mutation frequency than other relevant targets like the spike protein [53]. In the case that some or all of the cellular functions are mediated by a channel function of the E-protein the yeast complementation system can be developed together with similar systems in bacteria into versatile high-throughput screening systems for pore blockers.

## MATERIALS AND METHODS

### Cloning

A double-stranded DNA fragment encoding Ep-CoV2 (GenBank MN908947.3:26245-26472) for cloning into the various plasmids was purchased from IDT Inc. (Coralville, IA, USA).

For *in vitro* protein expression the Ep-CoV2 DNA fragment was amplified via PCR using primer pair 1 (table 1) and subsequently cloned into a modified pET24 vector (pET24Δlac) in which the lac-operator and the ribosome binding site (RBS) were replaced by the 5’-UTR of the *in vitro* expression vector pEXP-5-CT/TOPO (Invitrogen, Karlsbad, CA, USA). For cloning, pET24Δlac was first linearized with the restriction enzymes NdeI and SalI. The amplified Ep-CoV2 DNA fragment was then inserted into the vector backbone using NEBuilder® HiFi DNA Assembly (NEB, Ipswich, MA, USA). For Western blot analysis, Ep-CoV2 was N-terminally tagged via a TEV protease cleavage site (ENLYFQSYSFVS) with a FLAG epitope (DYKDDDDK). The DNA fragment encoding the desired FLAG-TEV tag was generated via PCR using primer pair 2 (table 1) and subsequently cloned into the NdeI restriction site of pET24Δlac/Ep-CoV2 using NEBuilder® HiFi DNA Assembly (NEB, Ipswich, MA, USA).

**Table 1:**
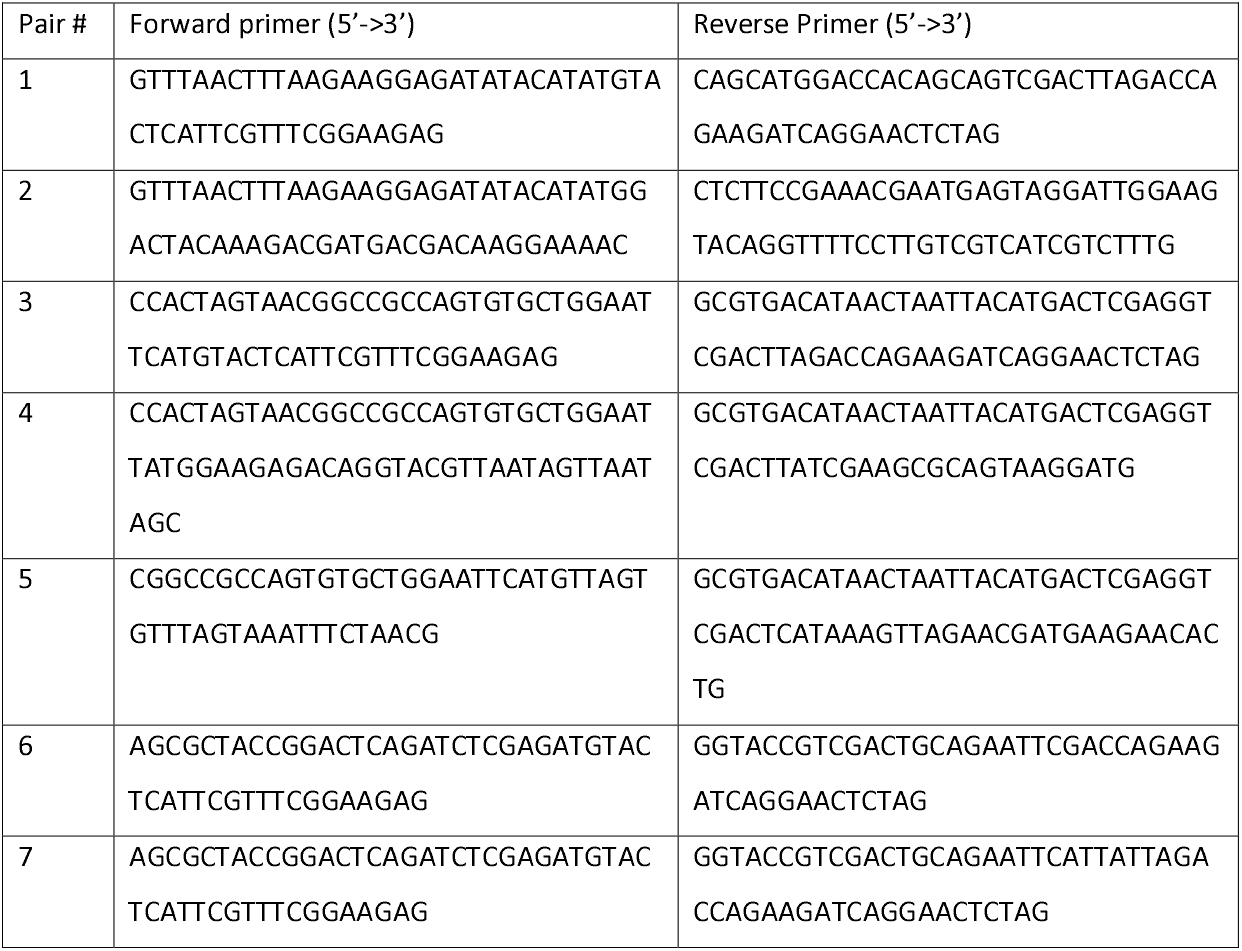
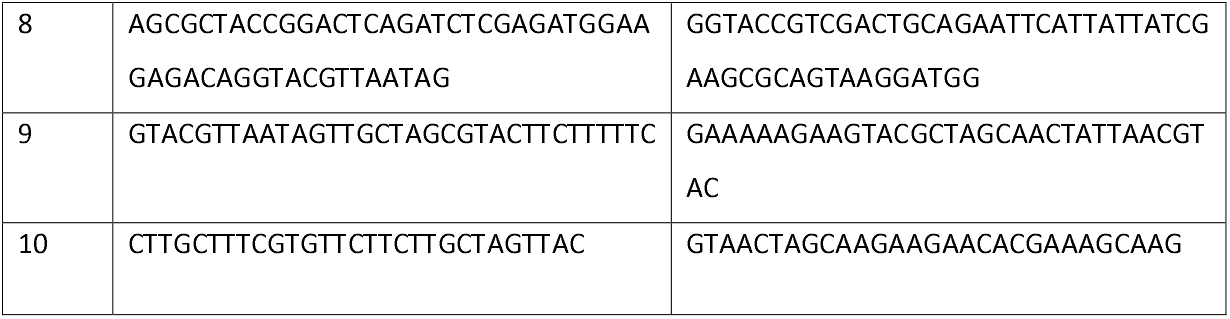
Sequences of used PCR primers.

For yeast complementation assays, the synthetic Ep-CoV2 DNA fragment was used as DNA template together with primer pair 3 or primer pair 4 (table 1) in a standard PCR in order to amplify the coding sequences of full length (Ep-CoV2) or truncated envelope protein (Ep-CoV2_™_), respectively, with appropriate overhangs. These fragments were subsequently inserted into a modified pYES2 shuttle vector (pYES2sh) in which the original P_*GAL1*_ promoter of pYES2 (Invitrogen, Karlsbad, CA, USA) was replaced by the methionine repressible P_*MET3*_ promoter. For cloning, pYES2sh was first linearized with the restriction enzymes NotI and EcoRI and amplified Ep-CoV2 or Ep-CoV2_™_ DNA fragments was inserted into the vector backbone using NEBuilder® HiFi DNA Assembly (NEB, Ipswich, MA, USA). The viral potassium channel Kcv_PBCV-1_ was used as a positive control in the yeast complementation assay [36]. For this purpose, the coding sequence of Kcv_PBCV-1_ was amplified with appropriate overhangs using primer pair 5 (table1) and subsequently cloned into pYES2sh as described above.

For expression in mammalian cells, Ep-CoV2 and Ep-CoV2_™_ were cloned into pEGFP-N2 which contains the eGFP sequence for generating C-terminal eGFP-tags. In order to generate the fusion protein Ep-CoV2::eGFP (Ep-CoV2_GFP_) the Ep-CoV2 DNA fragment was amplified via PCR using primer pair 6 (table 1) and subsequently cloned between the XhoI and EcoRI restriction sites using NEBuilder® HiFi DNA Assembly. In order to express Ep-CoV2 and Ep-CoV2_™_ without C-terminal eGFP-tag, the DNA fragments were amplified using primer pairs 7 and 8, respectively, thereby generating two stop codons and a reading frame shift downstream of the coding sequence.

The Ep-CoV2 mutants N15A and V25F were generated by site-directed mutagenesis as described in [54] using primer pairs 9 and 10, respectively.

Prior to further use of all generated constructs, their coding regions were checked by sequencing.

### Transfection of mammalian cells

For functional expression of Ep-CoV2, Ep-CoV2 mutants, Ep-CoV2_GFP_ and Ep-CoV2_™_, HEK293 and A549 cells were transfected 16 to 24 hours before the start of patch-clamp or fluorescence imaging experiments using TransfeX™ Transfection Reagent (LGC Standards GmbH, Wesel, Germany) according to the manufacturer’s instructions. HEK293 and A549 cells (German Collection of Microorganisms and cell cultures, Braunschweig, Germany) were grown at 37°C in a humidified 95% air/5% CO_2_ incubator in Dulbecco’s Modified Eagle Medium (DMEM; Gibco) supplemented with 10% v/v heat-inactivated fetal bovine serum, 100 U/ml penicillin G, 100 μg/ml streptomycin sulfate and 2 mM L-glutamine (all from Invitrogen). After reaching approximately 80% confluence mammalian cells were (co-)transfected in a 35 mm petri dish with 1 µg of the plasmid carrying the gene of interest and (if necessary) 1 µg of empty pEGFP-N2 or empty pIRES2-mRuby3 to enable identification of transfected cells via eGFP or mRuby3 fluorescence, respectively. pIRES2-mRuby3 was generated by replacing the sequence encoding for the green fluorescent protein eGFP in pIRES2-eGFP by a DNA sequence encoding for the red fluorescent protein mRuby3.

### Transformation of yeast cells

The K^+^ uptake deficient *S. cerevisiae* strains PLY240 (*MATa his3Δ200 leu2-3,112 trp1Δ901 ura3-52 suc2Δ9 trk1Δ51 trk2Δ50::lox-kanMX-lox*) [55] was transformed with the constructed pYES2sh plasmids using Frozen-EZ Yeast Transformation II kit (Zymo Research Europe GmbH, Freiburg, Germany) according to manufacturer’s instructions. PLY240 lacks the main K^+^-uptake systems *Trk1* and *Trk2* and is unable to grow in standard yeast media containing less than 10 mM potassium. Successfully transformed yeast cells were selected on SD-ura (20 g/l glucose, 6.9 g/l yeast nitrogen base without amino acids, 0.77 g/l drop-out -ura supplement, 20 g/l Agar-Agar) agar plates supplemented with 100 mM KCl. Cells were grown at 30 °C for 72 h and colonies were picked.

### Yeast complementation assay

For functional complementation assays, the transformed PLY240 strains were grown overnight in liquid SD-ura selection medium supplemented with 100 mM KCl. The following morning cells were harvested by centrifugation (500 x g, 4 min) and washed twice with ddH_2_O to remove residual potassium. Afterwards, the cells were suspended in ddH_2_O and the optical density at 600 nm (OD_600_) was adjusted to 1.0. Each well of a 24 well cell culture plate (Sarstedt, Germany) was loaded with 1 ml SD-ura-met medium supplemented with 0 to 100 mM KCl and then inoculated with 10 µL of the yeast cell suspension. In order to monitor cell growth, OD_600_ was measured every 30 min on a BioTek Epoch™ 2 Microplate Spectrophotometer (BioTek Instruments, Winooski, VT, USA) with constant double orbital shaking at 30°C. To determine the maximum growth rate (µ_max_), the growth curves were first smoothed by calculating the moving average (window width = 5 data points). Subsequently, the growth rate µ for two neighboring data points was determined by calculating the difference quotient

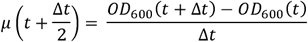

with t being the time and Δt the time between two neighboring data points. The highest value was then used as µ_max_.

### Patch-clamp experiments

On the day of the experiment, transfected HEK293 or A549 were separated by trypsinization, seeded at low density on 10*10 mm coverslips, and then incubated for 2 to 4 hours to allow adhering of cells on the glass surface. For patch-clamp experiments, coverslips were then transferred to a perfusion chamber filled with bath solution and placed on the stage of an inverted microscope. Transfected cells were identified by the fluorescence of co-expressed eGFP. Patch-clamp experiments were performed in the whole-cell configuration with an EPC-9 amplifier (HEKA Elektronik, Lambrecht, Germany) at room temperature (20-25°C). Patch-pipettes were pulled from borosilicate capillaries (DWK Life Sciences, Milville, NJ, USA) using a single stage glass microelectrode puller (PP-830, Narishige Group, Tokyo, Japan) resulting in pipettes with 2-3 MOhm resistances. The capillaries were coated at tapper with Sigmacote® (Merck KgaA, Darmstadt, Germany) and baked after pulling at 65°C for 45 min.

Whole-cell currents were generated in response to voltage steps from a holding potential of -80 mV to test potentials between -120 mV to +120 mV. Currents were filtered at 10 kHz using a low pass Bessel filter and sampled at 20 kHz without leak current subtraction. Membrane potentials were measured in current-clamp mode at zero current. Data was collected with PatchMaster (HEKA Elektronik, Lambrecht, Germany) and analyzed with FitMaster (HEKA Elektronik, Lambrecht, Germany). Liquid junction potentials (LJPs) were calculated using JPCalcWin (School of Medical Sciences, University of New South Wales, Sydney, Australia) and subtracted post recording. The standard pipette solution contained in mM: 10 NaCl, 130 potassium-D-gluconate, 1.93 CaCl_2_, 2 MgCl_2_, 5 4-(2-hydroxyethyl)-1-piperazineethanesulfonic acid (HEPES), 5 ethylene glycol-bis(β-aminoethyl ether)-N,N,N⍰,N⍰-tetraacetic acid (EGTA), and 2 MgATP. pH and osmolarity were adjusted to 7.2 with 1 M KOH and 298 mOsmol/kg with D-mannitol, respectively. The K^+^-free pipette solution contained cesium-D-gluconate instead of potassium-D-gluconate and pH was adjusted with 1 M CsOH. The standard bath solution contained in mM: 110 NaCl, 30 KCl, 1.8 CaCl_2_, 0.5 MgCl_2_, and 5 HEPES. For cesium experiments the bath solution contained in mM: 110 NaCl, 30 CsCl, 1.8 CaCl_2_, 0.5 MgCl_2_, 5 HEPES, and 10 mM tetraethylammonium chloride. Calcium-rich bath solution contained in mM: 110 NaCl, 20 CaCl_2_, 0.5 MgCl_2_, 5 HEPES, and 10 mM tetraethylammonium chloride. pH and osmolarity were adjusted to 7.4 with 1 M NaOH and 305 mOsmol/kg with D-mannitol, respectively. For cesium experiments with low external pH HEPES was replaced by 2-(N-morpholino)ethanesulfonic acid (MES) and pH was adjusted with 1 M HCl to 4.5.

### *In vitro* Protein expression, purification and Western-blot analysis

*In vitro* expression of Ep-CoV2 was done using the MembraneMax Protein Expression System (ThermoFisher Scientific, Waltham, MA, USA) following manufacturer’s instructions. In brief, 1 µg of template DNA (pET24Δlac/Ep-CoV2) was added to the transcription/translation reaction mixture in the presence of pre-assembled MSP1D1-His DMPC nanodiscs (Cube Biotech GmbH, Monheim, Germany). After initial incubation for 30 min at 37°C and 1250 rpm, feeding buffer was added and incubation was continued for additional 1.5 hours.

The reaction mixture was then added to a 0.2 mL HisPur Ni-NTA Spin column (ThermoFisher Scientific, Waltham, MA, USA), pre-equilibrated in PBS (phosphate buffered saline), adjusted to pH 7.4, with 10 mM imidazole, and incubated at constant shaking for 30 min at 37°C. Flow through was collected at 700 x g for 2 min, the column was then washed three times with 400 µL PBS adjusted to pH 7.4 with 25 mM imidazole, and finally proteins were eluted three times using 200 µL PBS adjusted to pH 7.4, with 250 mM imidazole. For western-blot analysis protein concentration was concentrated aprox. 30-fold using an Amicon Ultra-0.5 mL (Merck KgaA, Darmstadt, Germany) to a residual volume of 20 µL.

For western blotting, proteins were diluted in 4x LDS sample buffer (ThermoFisher Scientific) and separated using a Novex 4-12% Bis-Tris LDS Gel (ThermoFisher Scientific) in MES running buffer (ThermoFisher Scientific) at constant 150 V until the blue loading dye front reached the end of the gel. Proteins were transferred onto a Nitrocellulose membrane (ThermoFisher Scientific) using the iBlot2 dry blotting system (ThermoFisher Scientific). Membranes were blocked for 1 hour at RT using Pierce™ Clear Milk Blocking Buffer (ThermoFisher Scientific). Blots were incubated overnight at 4°C with mouse anti-FLAG tag mAB (Cell Signaling Technology, Danvers, MA, USA) at a final dilution of 1:1000 in blocking buffer. Blot were washed three times in Tris-buffered saline solution with 0.1% Tween for 5 min, incubated with Goat-anti mouse IRDye 680 mAB diluted 1:15,000 in blocking buffer for at least 1 hour at RT or overnight at 4°C and imaged using a LiCor Odyssey imaging system (LI-COR, Lincoln, NE, USA).

### Ion channel recordings

Recording of Ep-CoV2 mediated channel activity were done in horizontal suspended lipid bilayers by voltage clamp recordings. DPhPC (1,2-diphytanoyl-sn-glycero-3-phosphocholine) and DPhPS (1,2-diphytanoyl-sn-glycero-3-phospho-L-serine) were obtained from Avanti Polar Lipids (Alabaster, AL, USA) in chloroform solution. DPhPC and DPhPS were mixed 50/50 and 100 µL of the final solution was dried under a constant nitrogen stream and re-suspended in 100 µL n-decane to reach a final concentration of 25 mg/ml. Lipid bilayers were formed on the 50 µm cavity of a MECA4 chip (Ionera Technologies GmbH, Freiburg, Germany) using the pseudo-painting bubble technique resulting in stable lipid bilayer with a typical membrane capacitance of 40-60 pF. Nanodiscs loaded with Ep-CoV2 were added to the trans-side, i.e. the upper chamber with the reference Ag/AgCl electrode connected to ground on the HEKA EPC10-USB (HEKA Instruments) and voltage bias was applied at the Ag/AgCl electrode in the microcavity on the cis-side in reference to the ground electrode. After addition of 1-2 µL of the protein solution incorporation was typically monitored within 20-30 min. The trans-chamber was perfused with 10-20 sample chamber volumes of buffer to remove unbound proteins using a gravity driven perfusion system with vacuum suction. Unless stated otherwise, recordings were done in symmetrical buffer conditions using 250 mM KCl, 1 mM EGTA, and 10 mM HEPES, pH 7.4/KOH. In experiments with calcium the buffer solution in the trans-chamber was replaced with the respective Calcium buffer (125 mM CaCl_2_, 10 mM HEPES, pH adjusted to 7.4 with ⍰2.5 mM Ca(OH)_2_) or 125 mM CaCl_2_, 10 mM HEPES, pH adjusted to 7.4 with ⍰5 mM KOH) using the perfusion system. Data was low-pass filtered with a 10 kHz Bessel filter and sampled at 100 kHz. After digitization, data was additionally filtered to achieve satisfactory signal to noise ratio using a digital Bessel filter at 0.1 kHz.

### Microscopy

#### Single Molecule Detection/Total Internal Reflection Fluorescence (SMD/TIRF)

HEK293 cells transiently expressing Ep-CoV2_GFP_ were grown on coverslips (ø 25 mm). To favor strong adhesion of the cells to the glass, the coverslips were cleaned in a Zepto-B plasma furnace (Diener electronic GmbH, Ebhausen, Germany) and coated with a layer of 0.01% poly-D-lysine to favor strong adhesion of the cells to the glass. Plasma membrane patches were then isolated by an osmotic shock with ice cold distilled water as described previously [43]). The remaining plasma membrane patches on the glass coverslips were imaged on a Nikon Ti-E stand (Nikon, Konan, Minato-ku, Tokyo, Japan) with a CFI Apo TIRF 100x objective (NA 1.49, WD 0.12 mm). For TIRF imaging the focus in the back focal plane was moved off-center by controlling the position of a mirror with a single-axis micropositioner stage M-126. DG controlled by a C-863 Mercury Servo Controller (Physik Instrumente (PI), Karlsruhe, Germany). Plasma membrane patches and potential contaminations by remaining cortical ER were stained with red fluorescent CellMask™ Deep Red (CMDR) and ER-tracker™ Blue-white DPX (both from ThermoFisher Scientific), respectively. The fluorescent markers were excited/detected as follows: GFP (488 nm/500-550 nm), ER-Tracker™ (561 nm/577.5-646.5 nm) or CMDR (647 nm/ 662.5-799.5 nm).

#### Confocal laser scanning microscopy

Confocal laser scanning microscopy (CLSM) was performed on a Leica TCS SP or SP5 II system (Leica microsystems, Mannheim, Germany) equipped with 40 × 1.30 oil UV (HCX PL APO) objective. The external buffer for microscopy contained in mM: 140 NaCl, 4 KCl, 1 MgCl_2_, 5 D-mannitol, 10 HEPES, 2 CaCl_2_, pH 7.3 with an osmolarity of 310 mOsmol/kg. In all experiments mRuby3 fluorescence (561 nm/ 600-630 nm) was used as transfection control.

For monitoring the concentration of free Ca^2+^ in the cytosol ([Ca^2+^]_in_), HEK293 cells were loaded with the cell permeable Ca^2+^ sensor Fluo4-AM (ThermoFisher Scientific) for 30 min in microscopy buffer at a final concentration of 1 µM. In alternative experiments cells were loaded with FURA2-AM (ThermoFisher Scientific) as Ca^2+^ sensor by incubating cells for 30 min in buffer with 5 µM dye. Cytosolic pH was monitored by incubating HEK293 cells with the cell membrane permeable pH sensor BCECF-AM (2’,7’-Bis-(2-Carboxyethyl)-5-(and-6)-Carboxyfluorescein, Acetoxymethyl Ester) (Sigma Aldrich, St. Louis, MO, USA) for 30 min in microscopy buffer at a final concentration of 1.25 µM. The membrane potential of HEK293 cells was examined by loading the cells for 15 min with one of the two voltage sensitive dyes DiBAC_4_(3) (Bis-(1,3-Dibutylbarbituric Acid)Trimethine Oxonol) (AnaSpec, Fremont, CA, USA) or FluoVolt™ (ThermoFisher Scientific) diluted with microscopy buffer to a final concentration of 10 µM or 13.7 µM, respectively. All dyes were removed after cell staining by washing cells with dye free buffer.

All dyes were detected at 500-540 nm and excited at 360 nm (Fura2) or 488 nm (all other dyes). Images were taken at a resolution of 1024 × 1024 pixels and a scan speed of 200 or 400 Hz. Signals of positive controls were recorded in 5 s intervals for 15-30 min in total after adding the corresponding solution. The same procedures were employed with A549 cells.

## Acknowledgments

We thank Barbara Ehrlich, Andrew Marks and Edmond Buck for discussion, and Oliver Clarke and Francesca Vallese for providing anti-FLAG-tag antibodies.

## Author Contributions

AH, OR, GT, and AM conceptualized the study. TS, AH, SH, CH, RL, DT and OR collected and analyzed the data. GT, OR, AH and TS wrote the manuscript, TG searched literature and all authors provided edits and comments. AM, GT, AB and KS supervised the study.

