## Supplementary data for "SARS-CoV-2 envelope-protein corruption of homeostatic signaling mechanisms in mammalian cells"

### Figure S1

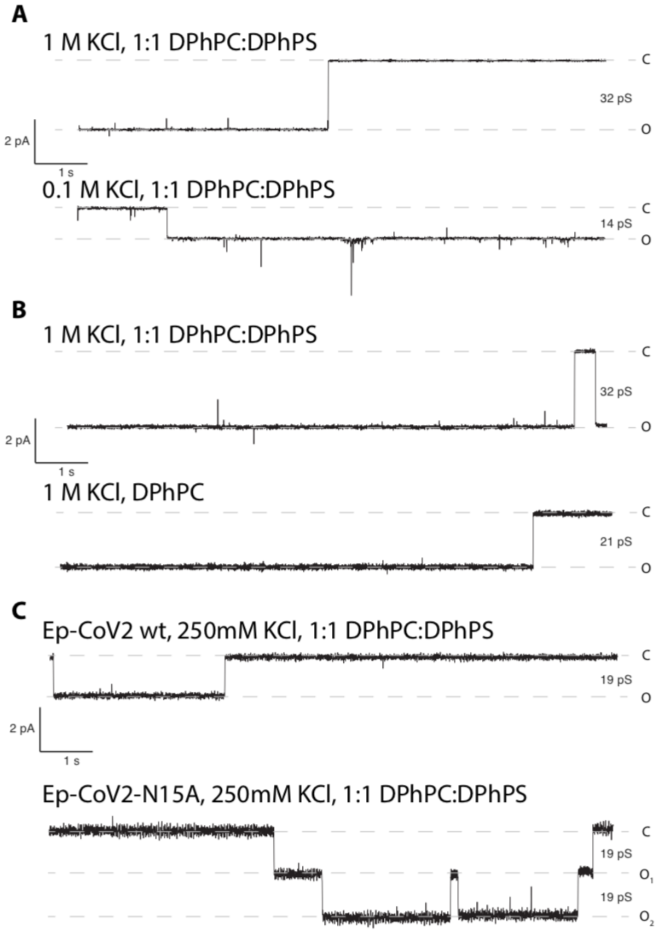


**Figure S1. Channel recordings of mutant Ep-CoV2 N15A and Ep-CoV2 wt under different experimental conditions. (A)** Effect of reducing KCl concentration in symmetrical recordings from 1 M (upper panel) to 0.1 M KCl (lower panel) on unitary conductance of Ep-CoV2. **(B)** Ep-CoV2 generated channel fluctuations in symmetrical 1 M KCl solution in absence and presence of anionic lipids. Unitary conductance in bilayer with 1:1 mix of neutral (DPhPC (1,2-diphytanoyl-sn-glycero-3-phosphocholine) and anionic lipid (DPhPS) (1,2-diphytanoyl-sn-glycero-3-phospho-L-Serin) (upper panel) and recording in neutral DPhPC-only bilayers (lower panel). **(C)** Channel fluctuations in symmetrical 250 mM KCl with mixed DPhPC:DPhPS (1:1) bilayer generated by Ep-CoV2 wt (upper panel) and E-CoV2 N15A (lower panel) channel. All recordings were done at -100 mV. Data was digitized at 100 kHz and digitally filtered for presentation at 100 Hz.

### Figure S2

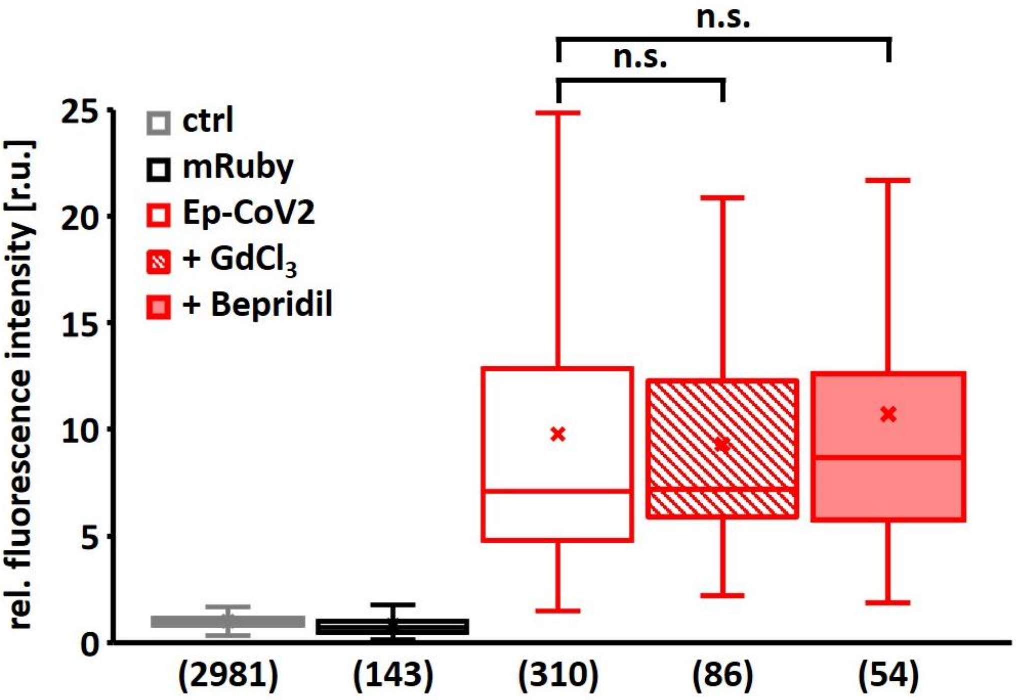


**Figure S2. $Ca^{2+}$ blockers do not abolish effects of Ep-CoV2 on $[Ca^{2+}]_{in}$.** Relative fluorescent intensities from $Ca^{2+}$-sensor Fluo4 in HEK293 cells expressing mRuby alone (black) or mRuby + Ep-CoV2 in absence (red, open bar), or presence of 10 µM $GdCl_3$ (red, dashed bar) or 20 µM Bepridil (red, solid bar). Data obtained as in Fig. 2B; data in absence of blockers are re-plotted from Fig. 2B. Number of measured cells in brackets. Boxes of box plots in A-D represent 25$^{th}$ and 75$^{th}$ percentiles; median is shown as horizontal line, arithmetic mean as cross. Bars indicate minimal and maximal value. Statistical significance was determined with unpaired two-tailed student t-test assuming unequal variances ($p > 0.05$, n.s.).

### Figure S3

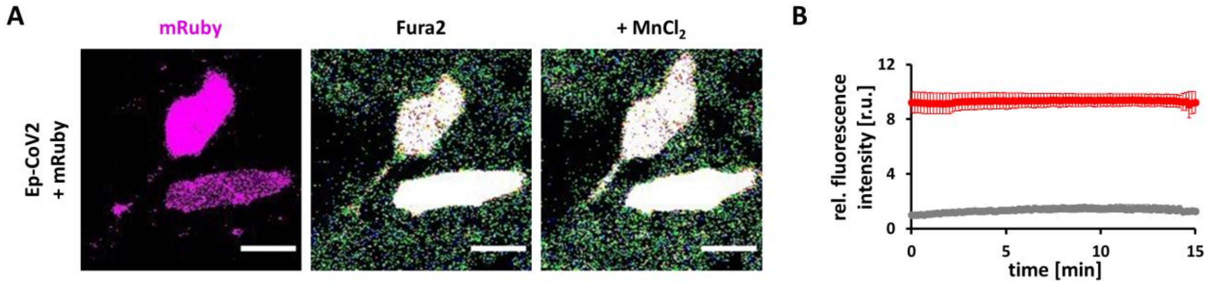


**Figure S3. Addition of $Mn^{2+}$ to external buffer does not quench elevated Fura2 signal in Ep-CoV2 expressing cells. (A)** Representative fluorescence images of HEK293 cells expressing mRuby + Ep-CoV2 with mRuby signal (left) and Fura2 signal in absence (central) and presence of 2 mM $Mn^{2+}$ in buffer (right). **(B)** Dynamics of mean Fura2-fluorescence intensity in un-transfected control cells (grey) and cells expressing mRuby + Ep-CoV2 (red) after addition of 2 mM $MnCl_2$ to external buffer at time zero. Mean ±SD from > 8 cells were normalized to fluorescence intensity of control cells prior to addition of $Mn^{2+}$. Scale bars = 20 µm.

### Figure S4.

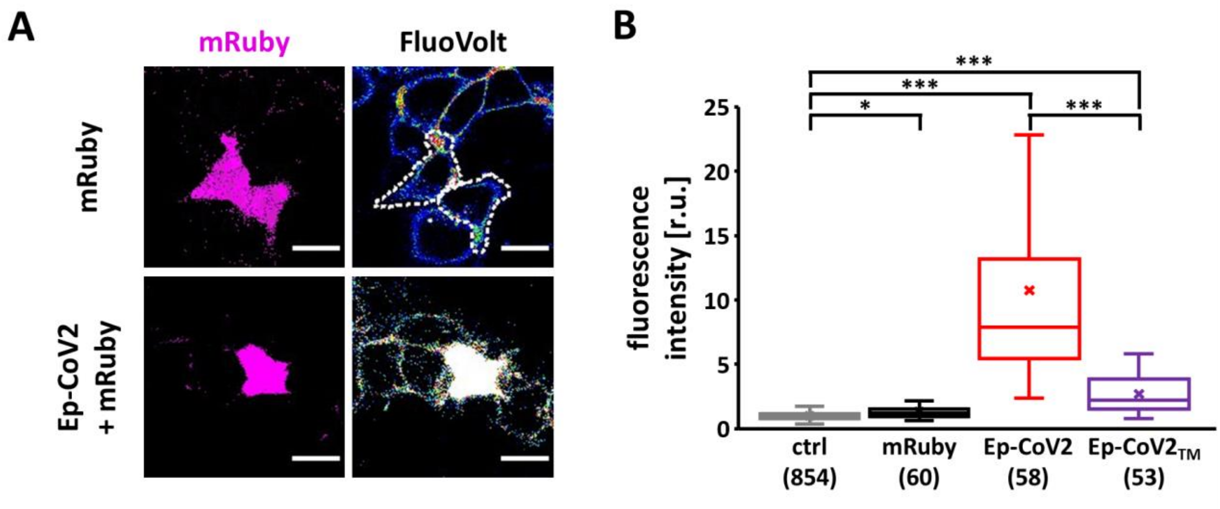


**Figure S4. Expression of Ep-CoV2 depolarizes plasma membrane. (A)** Representative fluorescence images of HEK293 cells expressing either mRuby alone (negative control; upper row) or mRuby + Ep-CoV2 (lower row). mRuby signal acts as a transfection control (first lane, magenta). Cells were loaded with voltage sensitive dye FluoVolt (second lane, 16 color LUT). Contours of mRuby positive cells in FluoVolt channel in upper row are highlighted with white dotted line. **(B)** Relative fluorescence intensity of FluoVolt in un-transfected control HEK293 cells (ctrl, grey) and HEK293 cells expressing mRuby alone (black), mRuby + Ep-CoV2 (red) and mRuby + Ep-CoV2$_{TM}$ (violet). Data are normalized to mean fluorescence intensity of un-transfected control cells; number of measured cells in brackets. Boxes of box plots in A-D represent 25$^{th}$ and 75$^{th}$ percentiles; median is shown as horizontal line, arithmetic mean as cross. Bars indicate minimal and maximal value. Statistical significance in (B) was determined with unpaired two-tailed student t-test assuming unequal variances ($p < 0.001$, ***; $0.001 < p < 0.01$, **; $0.01 < p < 0.05$, *; $p > 0.05$, n.s.). Scale bars 20 µm.
